# Benchmarking Molecular Dynamics Force Fields for All-Atom Simulations of Biological Condensates

**DOI:** 10.1101/2023.02.09.527891

**Authors:** Kumar Sarthak, David Winogradoff, Yingda Ge, Sua Myong, Aleksei Aksimentiev

## Abstract

Proteins containing intrinsically disordered regions are integral components of the cellular signaling pathways and common components of biological condensates. Point mutations in the protein sequence, genetic at birth or acquired through aging, can alter the properties of the condensates, marking the onset of neurodegenerative diseases such as ALS and dementia. While the all-atom molecular dynamics method can, in principle, elucidate the conformational changes that arise from point mutations, the applications of this method to protein condensate systems is conditioned upon the availability of molecular force fields that can accurately describe both structured and disordered regions of such proteins. Using the special-purpose Anton 2 supercomputer, we benchmarked the efficacy of nine presently available molecular force fields in describing the structure and dynamics of a Fused in sarcoma (FUS) protein. Five-microsecond simulations of the full-length FUS protein characterized the effect of the force field on the global conformation of the protein, self-interactions among its side chains, solvent accessible surface area and the diffusion constant. Using the results of dynamic light scattering as a benchmark for the FUS radius of gyration, we identified several force fields that produced FUS conformations within the experimental range. Next, we used these force fields to perform ten-microsecond simulations of two structured RNA binding domains of FUS bound to their respective RNA targets, finding the choice of the force field to affect stability of the RNA–FUS complex. Taken together, our data suggest that a combination of protein and RNA force fields sharing a common four-point water model provides an optimal description of proteins containing both disordered and structured regions and RNA–protein interactions. To make simulations of such systems available beyond the Anton 2 machines, we describe and validate implementation of the best performing force fields in a publicly available molecular dynamics program NAMD. Our NAMD implementation enables simulations of large (tens of millions of atoms) biological condensate systems and makes such simulations accessible to a broader scientific community.

## 1 INTRODUCTION

A protein’s ability to carry out its biological function has been traditionally attributed to its well-defined 3D structure, formed by deterministic folding of a polypeptide chain. ^1, 2^ The development of biochemical techniques beyond traditional crystallography, such as NMR, in tandem with advances in bioinformatics analysis and genome sequencing, revealed a class of proteins that do not fold into a unique 3D structure and are partially or completely disordered, the so-called intrinsically disordered proteins (IDPs) or proteins with intrinsically disordered regions (IDRs).^3^ These proteins typically have low sequence complexity, are rich in polar and charged hydrophilic residues and lack bulky hydrophobic groups that are known to form the core of structured proteins. The inherent “disorder” allows such proteins to access a plethora of conformations, ranging from extended coils to collapsed globules, thereby fulfilling multiple biological functions.^3, 4^ Despite lacking a well-defined 3D structure, an IDR can exhibit highly specific binding to its protein or nucleic acid target, a property used extensively for regulation of fundamental biological processes such as cell division, translation, and transcription, recruitment of proteins, and many others.^5^ For example, a mesh of disordered phenylalanine/glycine-rich nucleoporins (FG-Nups) regulates the passage of biomolecules in and out of a cellular nucleus. ^6^ The protein core of nucleosomes—the structural unit of chromatin— contains several disordered regions that recruit specific chromatin remodeling machinery depending on the pattern of post-translational modifications.^7^ Decoding the multifaceted structure-function relationship of proteins with IDRs holds key to molecular understanding of several biological processes and the origin of the associated diseases.

Over the last decade, several pioneering experimental studies have shown that proteins containing IDRs can spontaneously self-assemble into ”membrane-less” organelles to compartmentalize cellular functions and reactions.^8–10^ These organelles were found to exhibit properties of a liquid-like condensate ^11^ and to form or dissolve in the eukaryotic nucleus^12, 13^ or the cytoplasm^14, 15^ in response to biochemical signals. The aberrant phase separation processes have been linked to diseases associated with aging, such as amyotrophic lateral sclerosis (ALS) and frontotemporal dementia (FTD).^14, 16^ Recent experiment work elucidated how the amino acid composition of the IDPs affects the phase behavior and the physical properties of the condensates.^15, 17, 18^ However, precise molecular mechanisms governing the phase separation phenomena, in particular the effect of point mutations on the onset of neurodegenerative diseases, remain largely unknown.

Molecular simulations of the phase separation phenomena can provide microscopic insights inaccessible to experimental approaches.^19^ To date, the most successful simulations relied on so-called coarse-grained models, where one particle represents multiple atoms of the proteins (typically one amino acid) whereas the interactions between the particles derive from polymer physics principles or by matching experimental observables. Thus, an experimentally derived stickers-and-spacers model, where stickers account for protein-protein interactions and the spacers mediate contacts between stickers, was shown to accurately predict phase separation of prion-like disordered proteins.^20^ A residue-level hydrophobicity scale (HPS) model was parametrized against experimental data for several disordered proteins and used to chart the phase diagram of the DEAD-box helicase protein (LAF-1) and of the low complexity (LC) domain of fused in sarcoma (FUS) protein.^21^ Refined against experimental single-chain properties of 45 IDPs, the updated HPS model^22^ was used to elucidate the propensity of each amino acid to form condensates. Results of all-atom simulations were combined with experimental observations to build coarse-grained models, such as AWSEM-IDP,^23^ that retains molecular accuracy while offering sampling of large conformational space. However, a thermodynamic perturbation analysis has shown^23^ that the conformational space explored by IDPs is highly sensitive to even minor modifications of the underlying coarse-grained model.

As an alternative to coarse-grained approaches, all-atom molecular dynamics (MD) simulations can describe the equilibrium and kinetic properties of protein assemblies using molecular force fields that have been generated from the first principles.^24^ However, all-atom simulations of disordered proteins face multiple challenges, with the most critical ones being the accuracy of the molecular force-fields that govern the atomistic interactions and the sampling of a vast conformational space within the computationally accessible timescales. Conventional AMBER^25, 26^ and CHARMM^27, 28^ MD force-fields, which were developed to reproduce physical properties of folded proteins, are known to perform poorly when applied to IDPs, typically producing compact, globular conformations of too low of a radius of gyration to be experimentally feasible.^29, 30^ In the case of short intrinsically disordered peptides, enhanced sampling MD simulations ^31^ were able to reproduce an ensemble of conformations measured by experiment, although benchmark simulations have shown a pronounced dependence of the conformational ensemble on the molecular force field model.^32^ Recently, Pokorńa et. al. reported extensive all-atom MD simulations of protein-RNA interactions of a partial FUS construct containing both structured and disordered regions, describing rich conformational dynamics of the FUS–RNA system while also highlighting several irregularities arising from imperfections of the molecular force fields.^33^

Several groups have suggested improvements to the conventional force fields.^25, 27^ One of the most popular force fields for biomolecular simulations of folded proteins is AMBER ff14SB,^34^ which comes with the TIP3P^35^ water model and Joung-Cheatham ion parameters,^36^ which can be further augmented with non-bonded corrections^37^ for increased accuracy. Huang et. al.^38^ introduced CHARMM36m by modifying the CHARMM36 protein force field^28^ (which also comes with the TIP3P water model) using additional non-bonded fixes for modeling folded and disordered proteins. In the ff03ws model of Best and co-workers, the traditional TIP3P model of water^35^ was replaced with a newer, TIP4P model,^39^ which, upon modifications of the short-range interactions with proteins, provided accurate description of short IDPs, including in the crowded protein environment.^39, 40^ Izadi et. al.^41^ optimized the distribution of point charges on a water molecule, arriving with another four-point water model, OPC, that showed experimentally accurate molecular hydration free energies. The OPC water model, in conjunction with a more recent AMBER force-field, ff19SB,^42^ and Li-Merz^43^ ion parameters, is presently the recommended parameter set for biomolecular simulations using AMBER. ^44^ The use of the OPC water model was shown^45^ to noticeably improve the description of IDPs in comparison to equivalent simulations carried out using the TIP3P model.

In parallel to these efforts, Piana et al.^46^ introduced yet another four-point water model, TIP4P-D, that retained the TIP4P geometry but had its water dispersion coefficient (C_6_) increased by 50% in comparison to the TIP3P value.^35^ As the TIP4P-D model also retained the 12-6 Lennard Jones functional form, it offered compatibility with conventional biomolecular force fields. In particular, MD simulations of disordered proteins carried out using the ff99SB-ILDN^26^ protein force field, the TIP4P-D water and the CHARMM22 ion^27^ parameters showed expanded conformations with the radius of gyration in the range of NMR and SAXS predictions. To note, the ff99SB-ILDN force field includes an ILDN (Ile, Leu, Asp, Asn) torsion potential improvements of the conventional AMBER ff99SB^25^ force field, bringing the side-chain conformations in accordance with experiment.^26^ However, in the simulations of folded proteins, such as ubiquitin, the combination of ff99SB-ILDN and the TIP4P-D water was found to destabilize the protein’s native structure by *∼*2 kcal/mol in comparison to equivalent simulations carried out using the TIP3P water. ^46^ Best et. al. introduced backbone energy corrections for ff99SB to accurately balance the helix and coil populations in the simulations of unstructured peptides, ^47^ the corrections that are often used with the ILDN parameters^26^ and referred to as ff99SB*-ILDN. Finally, the DE Shaw lab introduced the a99SB-disp^48^ and DES-Amber^49^ force fields, both derived from the conventional ff99SB AMBER force field, to accurately model both structured and disordered regions in a protein. Also to note, both a99SB-disp and DES-Amber force fields use a modified TIP4P-D water model, optimized to work with each protein force field and correct the previously mentioned destabilization of structured proteins.

Here, we test nine MD force fields for their ability to describe the conformational dynamics of a full-length biological protein involved in the process of biological phase separation. As the subject of our test simulations, we use FUS, a 526-amino acid protein containing long IDRs, several key amino acids ubiquitous to phase separation and two structured regions, the RNA recognition motif (RRM) and the Zinc Finger Domain (ZnF).^50^ In addition to being an exemplary test system, the FUS protein is of great biological significance as it is involved in DNA repair, RNA recognition and binding and its condensate phase behavior is a marker of ALS and FTD. ^16, 51^ The majority of our test simulations were performed on the special purpose Anton 2 supercomputer,^52^ which allowed us to obtain relatively long (5 *µ*s duration) MD trajectory for each full-length FUS system and even longer (10 *µ*s duration) trajectories of RNA binding regions. Although the timescale of 5–10 *µ*s is not sufficient to fully sample the protein conformational landscape, it was found to be adequate to determine the average radius of gyration and to assess the propensity of a force field toward collapsed or extended conformations.^46^ As a primary benchmark of the force field accuracy, we used the radius of gyration of the FUS protein, which we experimentally measured using dynamic light scattering experiments. According to our benchmark simulations, the a99SB-disp forcefield^48^ appears to be an optimal choice for all-atom MD simulations of full-length biological condensate molecules. As this force field has not been available on supercomputer systems outside D.E. Shaw Anton, we describe our implementation of the a99SB-disp force-field for use with AMBER and NAMD simulation packages, and use our implementation for proof-of-principle all-atom MD simulation of a FUS/RNA condensate droplet.

## 2 MATERIALS AND METHODS

In this work, we performed equilibration simulations of a solvated model of a single full-length FUS protein using nine force-field combinations and two MD simulation engines, Anton 2^52^ and NAMD 2.14.^53^ We have also used Anton 2 to simulate two structured regions of the FUS protein (RRM and ZnF) in the presence of their respective RNA targets. Table 1 provides a concise summary of the runs performed.

**Table 1:** Summary of the benchmark runs. The first nine rows provide information about simulations carried out on the Anton 2 supercomputer. The next two rows provide information about the MD runs carried out using NAMD on TACC Frontera. The next row provides information on a coarse-grained simulation of a single FUS protein described in detail in Chou. et. al. ^57^ The last seven rows provide information about simulations of smaller RNA/protein systems carried out on Anton 2.

| Abbrev. | Protein force field | Water model | Ions force field | Simulation time and time step | ZnF bonds |
| --- | --- | --- | --- | --- | --- |
| ff14SB | ff14SB<sup>34</sup> + CUFIX<sup>37</sup> | TIP3P<sup>54</sup> | Joung-Cheatham<sup>36</sup> | 2 μs at 2.5 fs/step<br>3 μs at 4 fs/step | Yes |
| ff19SB | ff19SB<sup>42</sup> | OPC<sup>41</sup> | Li-Merz<sup>43</sup> | 5 μs at 2.5 fs/step | Yes |
| c36m | CHARMM36m<sup>38</sup> | TIP3P<sup>54</sup> | CHARMM36m<sup>55</sup> | 2 μs at 2.5 fs/step | No |
| ff03ws | ff03ws<sup>40</sup> | TIP4P (2005)<sup>39</sup> | Joung-Cheatham<sup>36</sup> | 2 μs at 2.5 fs/step<br>3 μs at 4 fs/step | Yes |
| tip4pd | ff14SB<sup>34</sup> + CUFIX<sup>37</sup> | TIP4P-D<sup>46</sup> | Joung-Cheatham<sup>36</sup> | 5 μs at 2.5 fs/step | No |
| desam | DES-Amber-SF1.0<sup>49</sup> | TIP4P-D<sup>46</sup> | DES-Amber<sup>49</sup> | 5 μs at 2.5 fs/step | No |
| desam0.9 | DES-Amber-SF0.9<sup>49</sup> | TIP4P-D<sup>46</sup> | DES-Amber<sup>49</sup> | 5 μs at 2.5 fs/step | No |
| aastar | ff99SB-star-ILDN<sup>26,47</sup> | TIP4P-D<sup>46,48</sup> | CHARMM22<sup>27</sup> | 5 μs at 2.5 fs/step | No |
| aadisp | a99SB-disp<sup>48</sup> | TIP4P-D-1.6<sup>48</sup> | CHARMM22<sup>27</sup> | 5 μs at 2.5 fs/step | No |
| aadispNAMD | a99SB-disp-NAMD | TIP4P-D-1.6 | CHARMM22 | 5 μs @ 2 fs/step | No |
| aastarNAMD | ff99SB-star-ILDN | TIP4P-D | CHARMM22 | 5 μs @ 2 fs/step | No |
| c36mNAMD | CHARMM36m | TIP3P | CHARMM36m | 1 μs @ 2 fs/step<br>+4 μs @ 4 fs/step | No |
| CG-KH | Kim-Hummer<sup>56</sup> | Implicit solvent | | 4 μs at 10 fs/step | Yes<sup>57</sup> |
| RRM-ff14SB | ff14SB<sup>34</sup> + CUFIX<sup>37</sup> | TIP3P<sup>54</sup> | Joung-Cheatham<sup>36</sup> | 10 μs at 2.5 fs/step | No |
| RRM-tip4pd | ff14SB<sup>34</sup> + CUFIX<sup>37</sup> | TIP4P-D<sup>46</sup> | Joung-Cheatham<sup>36</sup> | 10 μs at 2.5 fs/step | No |
| RRM-aastar | ff99SB-star-ILDN<sup>26,47</sup> | TIP4P-D<sup>46</sup> | CHARMM22 | 10 μs at 2.5 fs/step | No |
| RRM-aadisp | a99SB-disp<sup>48</sup> | TIP4P-D-1.6<sup>48</sup> | CHARMM22 | 10 μs at 2.5 fs/step | No |
| ZnF-tip4pd | ff14SB<sup>34</sup> + CUFIX<sup>37</sup> | TIP4P-D<sup>46</sup> | Joung-Cheatham<sup>36</sup> | 10 μs at 2.5 fs/step | Yes |
| ZnF-aastar | ff99SB-star-ILDN<sup>26,47</sup> | TIP4P-D<sup>46</sup> | CHARMM22 | 10 μs at 2.5 fs/step | Yes |
| ZnF-aadisp | a99SB-disp<sup>48</sup> | TIP4P-D-1.6<sup>48</sup> | CHARMM22 | 10 μs at 2.5 fs/step | Yes |

### 2.1 General MD protocols

Initial equilibration simulations were performed using the classical MD package NAMD,^53^ periodic boundary conditions, and a 2 fs integration time step. The ff14SB force field was used to describe proteins, TIP3P for water, and ions were described using Joung-Cheatham parameters. RATTLE^58^ and SETTLE^59^ algorithms were applied to covalent bonds that involved hydrogen atoms in protein and water molecules, respectively. The particle mesh Ewald (PME) ^60^ algorithm was adopted to evaluate the long-range electrostatic interaction over a 1 Å-spaced grid. Van der Waals interactions were evaluated using a smooth 10-12 Å cutoff. Multiple time stepping was used to calculate local interactions every time step and full electrostatics every three time steps. The Nose-Hoover Langevin piston pressure control^61, 62^ was used to maintain the pressure of the system at 1 atm by adjusting the system’s dimension. Langevin thermostat^63^ was applied to all the heavy atoms of the system with a damping coefficient of 0.1 ps*^−^*^1^ to maintain the system temperature at 310 K. Production runs in the NPT ensemble were completed on the D. E. Shaw Research Anton 2 supercomputer.^52^ The simulations on Anton 2 employed a set of parameters equivalent to those listed above, except for the use of the Martyna–Tobias–Klein barostat, the Nose-Hoover thermostat and the k-space Gaussian split Ewald method for calculations of the electrostatic interactions.

### 2.2 Preparation of full-length FUS systems

The initial conformation of the FUS protein was derived from an equilibrated CG simulation of a single FUS protein at 310 K, as reported previously.^57^ The protein was described at one-bead-per-amino acid resolution using the Kim-Hummer model^56^ for amino acid interactions and simulated for 4 *µ*s. The equilibrium CG structure was selected to be representative of the average radius of gyration observed within the 4 *µ*s simulation. To back-map the disordered fragments of the CG model to an all-atom representation, we replaced each bead with an amino acid, maintaining the CG backbone orientation but shrunk to 10% of it’s original size for non-aromatic side-chains and 25% of it’s original size for aromatic and proline side-chains, while maintaining the side-chain orientation. For the two structured regions of FUS, the RNA recognition motif (RRM, residues 278 to 385) and the Zinc Finger domain (ZnF, residues 419 to 454), we used the crystal structure coordinates (protein data bank entires 6GBM and 6G99, respectively^50^), which replaced the corresponding structured parts of the CG model. The structured regions were also shrunk in a similar fashion as the disordered regions, and the full-length protein was then subjected to minimization in vacuum using the NAMD simulation package, allowing the amino acids to attain their target size while avoiding development of steric clashes and ring piercings within the amino acid side chains. The amino acids in the structured regions grew back to the pdb structures, avoiding clashes, with heavy atom root mean square deviation of 0.73 Å and 0.86 Å for RRM and ZnF respectively. The FUS protein was then solvated in a cubic volume of water and ionized to produce a 150 mM KCl solution.

The solvated system was parametrized within the standard ff14SB force-field model^34^ using the *tleap* module of AmberTools16.^64^ We added four physical bonds between Zn and SG atom of cysteine to maintain the Zn^2+^ atom at the center of the four cysteine residues in the ZnF domain, also using the tleap tool, as we were unable to enforce such restraints as extrabonds on Anton 2. The bond, angle and dihedral parameters describing the chemical bond between the zinc atom and the SG atom of a cysteine residue were taken from the equivalent ff14SB parameters^34^ of the disulfide bonds in a cysteine residue. The potassium and chloride ions in the system were described using the Joung-Cheatham monovalent ion parameters^36^ whereas the Zn^2+^ atom was described using the Li-Merz parameters for highly charged metal ions.^65^ CUFIX corrections to non-bonded interactions between specific pars of charged chemical moieties were applied.^37^ The final system was a cubic box measuring *∼*150 Å on each side and contained *∼*252,000 atoms, Fig. 1a. This system was used as the initial structural model for the preparation of the ff14SB system (Table 1). The same protocol was followed in an exact fashion to create three more systems as described below with a few modifications.

**Figure 1:**
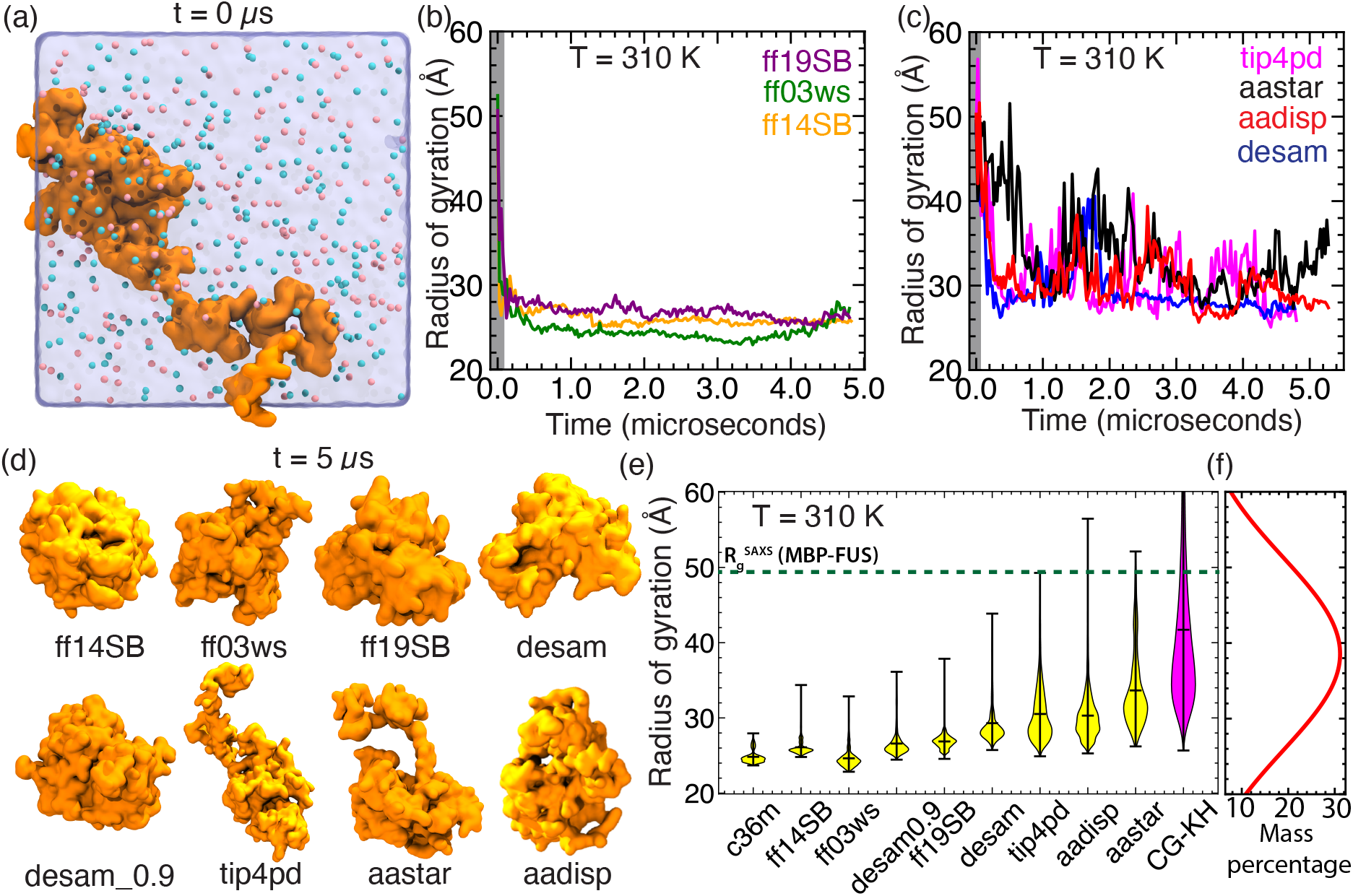
Radius of gyration of a monomeric FUS protein across MD force fields. (a) Initial configuration of the simulation system. The FUS protein is shown as an orange molecular surface, the volume occupied by the solvent as as a blue translucent surface and potassium and chloride ions as red and blue spheres, respectively. (b,c) The radius of gyration of the FUS protein as a function of simulation time for the seven parameter sets. The abbreviated names of the parameter sets are defined in Table 1. (d) Instantaneous configurations of the FUS protein at the end of the 5 *µ*s MD runs. (e) Violin plots of the radius of gyration values explored by the FUS protein in the corresponding MD runs. The first 50 ns of each trajectory (gray rectangles in panels b and c) were omitted from the analysis. The c36m *R*_g_ values were taken from a 2 *µ*s simulation. The Kim-Hummer (CG-KH) *R*_g_ values were taken from a previously reported ^57^ 4 *µ*s coarse-grained simulation. Dashed line denotes the average *R*_g_ value of a FUS protein tagged with a maltose binding protein measured by SAXS experiments. ^70^ The range of *R*_g_ values obtained from DLS experiments is shown as an orange rectangle. (f) Quantification of a single FUS particle size distribution in terms of the *R*_g_ as determined by dynamic light scattering at 60 min incubation time for 1 *µ*M FUS. The result shown is a gaussian fit over the raw data which is shown in SI Fig. S2.

A second system was created without having explicit physical bonds between the Zn ion and the cysteine residues and used for the preparation of the c36m,^55^ tip4pd,^46^ desam,^49^ desam0.9,^49^ aastar^26, 47^ and aadisp^48^ systems (Table 1). A third system was created by replacing the ff14SB force field in the *tleap* script with the force field files for the ff03w and TIP4P (2005) models obtained from Best. et. al.^40^ The ff03w force field was converted to ff03ws using a custom python script from Best et. al.,^40^ with the final system being the starting point for the ff03ws system (Table 1). A fourth and final system was created by replacing all force-fields in the tleap script with the ff19SB force field^42^ for proteins, OPC model^41^ for water and Li-Merz parameters^43^ for ions. The force field leaprc files and the *tleap* module for this specific case, was used from AmberTools21.^44^ To note, the last two systems kept the physical bonds between the zinc and cysteine SG atoms, in the ZnF domain. Finally, harmonic restraints were applied to the C*_α_* atoms of the full-length protein, using the starting structure as reference, and all four systems were simulated in the NPT ensemble using NAMD *∼*10 ns, which was sufficient for the system to attain its equilibrium volume.

The equilibrated systems were prepared for Anton 2 runs by converting the AMBER input format to the Anton dms format using the viparr-convert-prmtop module on Anton 2. To note, the AMBER restart files (rst7) generated from NAMD could not be directly used with the viparr-convert-prmtop module as they lacked information about atom velocities. We used custom tcl/python scripts to import atom velocities from the NAMD velocity restart files, adding them to the end of the restart rst7 file in the proper AMBER format. The dms files for the systems already built with the target force-field (ff14SB,^34^ ff03ws,^40^ ff19SB^42^) were run as is, upon using the viparr-build-constraints module to add the SHAKE constraints. The other force-field models (c36m,^55^ tip4pd,^46^ desam,^49^ desam0.9,^49^ aastar,^26, 47^ and aadisp^48^) were prepared starting from the dms file generated from their equilibrated AMBER input files and replacing the force-field using the viparr module of Anton 2. Because of the limitations of the viparr module in parametrizing non-standard residues, we were not able to add physical bonds for the zinc atom in the ZnF domain for the following five simulation systems: c36m, desam, desam0.9, aastar and aadisp. All models except c36m were each simulated for 5 *µ*s on Anton 2. For two systems, ff14SB and ff03ws, we applied hydrogen mass repartitioning (HMR) after 2 *µ*s to increase simulation time step to 4 fs using a custom implementation from Shabane et. al., ^45^ which is also one of the two suggested approaches in the Anton documentation. As also mentioned in the documentation, HMR has not been extensively used on Anton 2, which is why we did not use HMR for the majority of our simulations.

Two additional systems were generated using either the standard CHARMM36m^55^ (c36m) force field or our custom implementation of the aadisp force fields (described below). The two models were simulated for 5 *µ*s using NAMD on Frontera of the Texas Advanced Computing Center. After 1 *µ*s, the c36m NAMD simulation was run with a 4 fs time step using HMR.

### 2.3 Preparation of a99SB-disp systems for NAMD runs

Starting from a CHARMM-format PDB of one FUS protein dissolved in water and ions, we built AMBER-format files employing the following force fields: a99SB-disp ^48^ for protein, CHARMM22^27^ for ions, and the TIP4P-D-1.6^48^ four-point water model. We first built non-hydrogen atom PDBs of the separate components of the system (protein, ions, water), the residue and atom names of which were modified from CHARMM to AMBER nomenclature. The *tleap* AmberTool was then used to generate three separate library files using custom lib, frcmod and leaprc files (available as Supporting Information). To note, the existing TIP4P-EW water model was redefined to be TIP4P-D-1.6, i.e. with the properties described in Ref. 48. Another *tleap* script was used to combine those three lib files into an AMBER-format parameter/topology file (prmtop) and coordinate file (rst7). The combined prmtop file was modified to add extra backbone dihedral angles to the protein using the *ParmEd* python module. Following that, the rst7 coordinate file was centered about the origin. The modified prmtop and rst7 files were then used as input for NAMD simulations.

After 1 ns equilibration performed having the protein non-hydrogen atoms harmonically restrained to their initial coordinates with a force constant of 0.5 kcal mol*^−^*^1^ Å*^−^*^1^, NAMD simulations closely followed the “General MD protocols” set forth above. The pressure *P* was maintained at 1 atm using the Nośe-Hoover barostat, the temperature *T* held constant at 310 K using the Langevin thermostat. The resulting 5 *µ*s NAMD trajectory of a FUS protein at 310 K was analyzed alongside the trajectory from Anton 2, since they were different implementations of the same force field.

### 2.4 Preparation of RRM-RNA and ZnF-RNA systems

The initial coordinates of the RRM and ZnF domains bound to RNA were taken from their respective NMR structures, protein data bank entries 6GBM and 6G99.^50^ The RRM domain was resolved in a complex with a 23-residue stem loop RNA whereas the ZnF domain was resolved in a complex with a 5-residue UGGUG ssRNA. Each system was placed in a 150 mM KCl solution and parametrized to run with the Amber ff14SB^34^ protein force field, TIP3P^54^ water, Joung-Cheatham^36^ parameters for ions and Amber OL3^66^ parameters for the RNA and CUFIX^37^ corrections for non-bonded interactions implemented using *tleap*. The final RRM-RNA system was a cube *∼*65 Å on each side and contained *∼*28,000 atoms. Similarly, the ZnF-RNA system measured *∼*50 Å on each side and contained *∼*12,500 atoms. Both systems were equilibrated in the NPT ensemble using NAMD for *∼*10 ns while restraining the backbone atoms of each protein-RNA complex about their experimental coordinates. The zinc atom and the four cysteine SG atoms in the ZnF-RNA system were also restrained for the NAMD equilibration. Similar to the full-length FUS system, the equilibrated systems were converted to Anton 2 dms format, while replacing the force fields to create seven systems as specified in Table 1 (see the last seven rows). For the Anton simulations of the ZnF-RNA system, the zinc atom and the four SG atoms of the nearby cysteines were restrained to their initial coordinated to maintain the structure of the zinc atom coordination site. All seven systems were simulated using Anton 2 as described above for 10 *µ*s each.

Separately, two Amber-based versions of the RRM-RNA system were built with the *tleap* AmberTool for the purpose of running NAMD simulations. Starting from the fully solvated and ionized system described above, we generated systems with TIP4P-D-1.6^48^ four-point water, CHARMM22^27^ ions, modified AMBER ff14^67^ for ssRNA, and for protein either (1) a99SB-disp,^48^ or (2) a99SB-star-ILDN.^26, 47^ All-atom MD simulations of these two systems were performed with NAMD, starting from identical initial coordinates. The C*_α_* and C1*^!^* atoms were restrained for the first 10 ns, which was followed by unrestrained equilibration in the NPT ensemble for up to 10 *µ*s.

### 2.5 Contact analysis

To quantify the change in contacts in the protein-nucleic acid systems, we calculated the number of native contacts, Q, as^68^

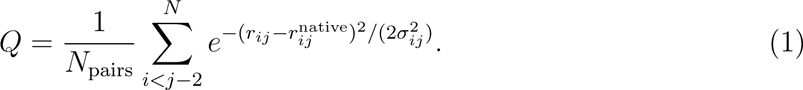

In this equation, *N*_pairs_ is the number of pairs of residues that are in contact in the PDB structure, *r_ij_* is the instataneous distance between the C*_α_* atoms of residues *i* and *j*, 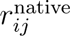 is the same distance in the PDB structure, and *σ_ij_* is (1 + *|i − j|*^0.15^). A contact is defined if a non-hydrogen atom of a protein residue is within 3 Å of a non-hydrogen atom of the nucleic acid residue.

### 2.6 Dynamic light scattering (DLS) measurements

Following a protocol described earlier,^69^ we performed DLS measurements using 500 nM, 1 *µ*M and 5 *µ*M solution of wild type FUS protein under the following buffer conditions: 1 M KCl, 1 M Urea, 1.5 mM *β*ME, 5% glycerol, 50 mM Tris-HCl and pH of 7.4. All the components of the solution mixture went through a 0.22 *µ*M syringe filter before mixing. The samples were held in a non-binding 96-well plate (Corning, Product Number 3881) during the measurements. The results of the DLS measurements were acquired using the Wyatt DynaPro Plate Reader II (Wyatt Technology) and transformed into size distribution using the Dynamic 7 software (Wyatt Technology). After extracting the distributions into a set of .csv files, a MATLAB script was applied to visualize the data.

## 3 RESULTS

To evaluate performance of MD force fields in explicit solvent simulations of biological condensates, we used Anton 2 supercomputer to run multiple simulations of a full-length FUS protein submerged in electrolyte solutions and also of two smaller systems each containing an RNA molecule specifically bound to a structural domain of FUS at the onset of each simulation. Our Anton 2 simulations explored nine force-field combinations, Table 1. The resulting trajectories were analyzed to characterize the conformational ensemble explored by the FUS protein, stability of its structured domains, the interactions of the structured domains with RNA, and the kinetics of the structural fluctuations. We implemented best performing force fields in NAMD2 and tested our implementation by repeating simulations of the full-length protein and the two RNA systems. Materials and Methods section provides detailed description of the simulation procedures. Supplementary Information provides stepby-step guide to using best performing force fields in NAMD and examples of the force field conversion workflow.

### 3.1 Global conformation ensemble of a full-length FUS protein

As detailed in Methods, we built an all-atom model of a full-length FUS protein submerged in a cubic volume of 150 mM KCl solution, Fig. 1a, and simulated the model using nine force field combinations for 5 *µ*s. Over the course of these free equilibration simulations, the FUS protein was observed to undergo major conformational changes, which we characterized by computing the protein’s radius of gyration and its end-to-end distance.

With the majority of the FUS protein being intrinsically disordered, its radius of gyration, *R*_g_, presents a convenient measure of the protein’s overall shape. The starting conformation of FUS that we chose for all our benchmark runs has an *R*_g_ value of approximately 50 Å and an asymmetric, elongated shape, Fig. 1a. Within the first 100 ns of the simulation carried out using a widely used parameter set, ff14SB for protein and TIP3P for water, the FUS protein spontaneously collapsed into a compact, symmetric globule of *R*_g_ value below 30 Å, Fig. 1b. Upon collapse, the conformational dynamics of the protein was subdued and the protein retained its compact conformation of 25 Å average *R*_g_ value for the remainder of the 5 *µ*s simulation. Similar collapse was observed in a 2 *µ*s CHARMM36m run (SI Fig. S1) and also in both 5 *µ*s ff03ws and ff19SB runs performed using the TIP4P (2005) and OPC water models, respectively. Qualitatively different behavior was observed in the simulations that employed the TIP4P-D water model, Fig. 1c. Although initial collapse was observed for some parameter sets, the collapse was reversible and the protein was able to explore relatively high *R*_g_ values over the course of the 5 *µ*s simulations. Figure 1d shows instantaneous configurations of the FUS protein at the end of the benchmark runs.

To quantitatively assess the outcome of the benchmark runs, we plotted the distributions of the *R*_g_ values adopted by the protein in all runs, Fig. 1e. The protein force-field combinations that have the lowest *R*_g_ are ff14SB and c36m (average *R*_g_ of *∼*25 Å), followed by ff19SB (average R_g_ of *∼*27 Å), which is not surprising as they were parametrized to reproduce the properties of folded proteins. Unexpectedly, a similar range of *R*_g_ values was also observed in the simulation that employed a parameter set specifically developed to describe disordered proteins, ff03ws. Slightly higher average *R*_g_ values were observed for the two amber force field variants, desam-0.9 and desam, used in combination with the TIP4P-D model, with the average *R*_g_ for the latter being *∼*30 Å. Unexpectedly, an even broader distribution of the *R*_g_ values was observed when an older amber force field, ff14SB, was combined with a TIP4P-D model and CUFIX corrections, a parameter set referred here as tip4pd. Finally, the highest average *R*_g_ values, *∼*34 Å and *∼*30 Å, were recorded for the aastar and aadisp parameter sets, respectively, with instantaneous *R*_g_ values covering a broad range, from 26 to 54 Å, in the case of aadisp.

For reference, we plot in Fig. 1e the distribution of *R*_g_ values observed in a 4 *µ*s CG simulation of a single FUS protein^57^ carried out using the Kim-Hummer model. ^56^ With the average *R*_g_ value of *∼*44 Å and a broad range of conformational fluctuations, the CG model predicts considerably higher *R*_g_ values than any all-atom model. Experimentally, the *R*_g_ value of 48.7 Å was measured by SAXS for single FUS proteins tagged with a 370-residue maltose binding protein (MBP). At the same time, the *R*_g_ of FUS in a globular conformation was estimated from its molecular weight to be 30.2 Å.^70^ Our own dynamic light scattering measurement, Fig. 1f, (see Materials and Methods) suggest a 30-to-50-Å range of *R*_g_ values, which compares favorably with the ensemble sampled by CG KH model and overlaps with the conformational ensemble sampled by tip4pd, aastar and aadisp models. Additional experiments confirmed that our DLS measurements characterize the conformations of FUS monomers because the size distribution remained unchanged when increasing the concentration of FUS or the duration of the measurement, SI Fig. S2 and S3. All other models appear to be sampling FUS conformations that correspond to a globular state.

Another measure of global protein conformation is the distance between the first and last residue of the protein, *i.e.*, the end-to-end distance. The end-to-end distance of biomolecules is experimentally accessible by means of single molecule Förster Resonance Energy Transfer (smFRET) measurements, which, in the case of FUS, was used to characterize association between FUS variants and FUS–RNA interactions.^51, 71^ In our benchmark simulations, a constant end-to-end distance of *∼*30 Å was observed for the ff14SB parameter set, Fig. 2a, indicating a frozen globular-like state. For the c36m parameter set, the end-to-end distance was observed to vary within a narrow, 10 and 40 Å range (SI Fig. S4). A higher variance of the end-to-end distance was seen for the ff03ws and ff19SB parameter sets, ranging from 20 to 90 Å for ff03ws, Fig. 2a. Qualitatively different end-to-end dynamics were observed for the models that employed the TIP4P-D water model, Fig. 2b, characterized by frequent large amplitude fluctuations, the overall range of values (10 to 180 Å) and considerably higher average values. Figure 2c characterizes the distribution of the end-to-end distances for all molecular force field models used in our benchmark runs.

**Figure 2:**
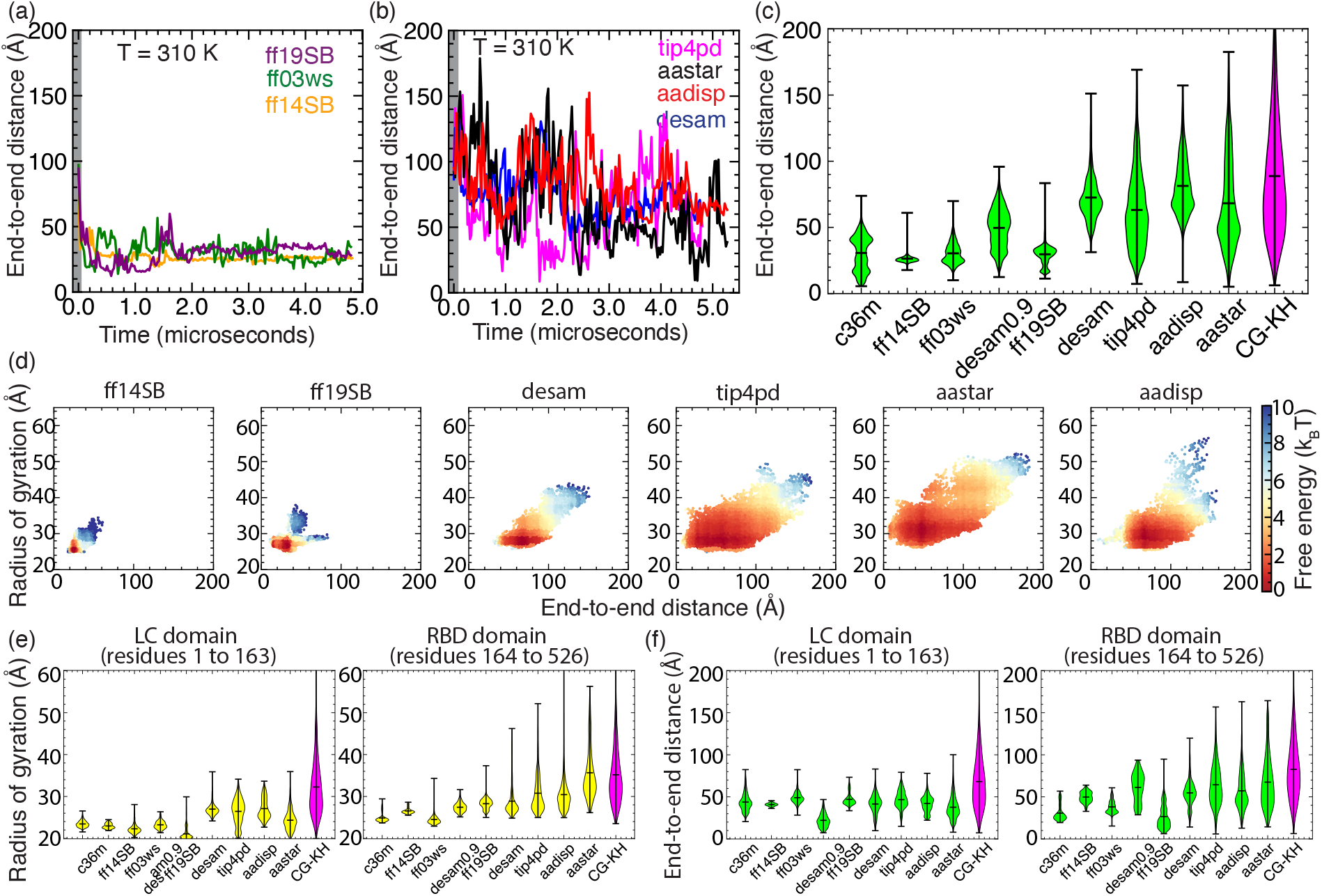
End-to-end distance and relative free energy of FUS conformations. (a,b) The end-to-end distance of the FUS protein as a function of simulation time for the six parameter sets. (c) Violin plot of the FUS end-to-end distance explored in the respective MD simulations. The first 50 ns of each trajectory (gray rectangles in panels a and b) were omitted from the analysis. (d) The free energy map of the FUS protein’s conformation as a function of its radius of gyration and end-to-end distance. Each map was constructed by Boltzmann inversion of the conformations sampled by the protein within 5 *µ*s trajectory, omitting the first 50 ns. The sampling rate was 0.24 ns; the bin size along both coordinates was 1 Å. The reference free energy state for all maps is set at 0, corresponding to a maximum theoretical probability of 1. Given two points on a map, the difference in free energies represents the likelihood of going from one state to the other, favored in the direction of the lower value. (e) Violin plot of the radius of gyration of the LC domain (left) and the RBD domain (right) of the FUS protein, explored in the respective MD simulations. (f) Same as (e) but for the end-to-end distance of the LC (left) and RBD (right) domains of FUS.

Having sampled both the *R*_g_ and the end-to-end distance distributions, we computed the relative free energy maps of single FUS conformations for all the parameter sets, Fig. 2d and SI Fig. S5. For these calculations, the free energy of a FUS conformation was defined as the Boltzmann probability of finding a protein within the specified range of *R*_g_ and end-to-end distance values. The maps indicate that the simulations performed using either ff14SB, ff19SB or ff03ws parameter set explored a much smaller conformational space than equivalent simulations that employed the TIP4P-D model, i.e., tip4pd, aastar and aadisp. For the aastar and aadisp force fields, the major fraction of easily switchable conformations (*<* 2*k*_B_T) correspond to an *R*_g_ (y-axis) values between 25 to 35 Å, which are closer to the globular state. Nevertheless, if compared to ff14SB, these low free energy FUS conformations are not static, fluctuating rapidly, as the end-to-end distance changes within 10-to-140-Å range. The access to extended states (outliers in the free energy plot), which are more frequently observed in the CG simulations, is associated with a free-energy penalty of about 4–8 *k*_B_T. Thus, the aastar and aadisp force fields provide access to extended FUS conformations albeit at low frequency, which leaves some room for future improvement and computational exploration using enhanced sampling methods.

The N-terminal of FUS contains a low complexity domain (LC, residues 1 to 163) which, while being completely disordered, is known to engage into biologically important interactions with client proteins and facilitate self-assembly *via* liquid-liquid phase separation (LLPS).^72^ The C-terminal of FUS contains the RNA binding domain (RBD, residues 164 to 526)—two structured regions interspersed with IDRs, which is primarily known for RNA recognition and binding.^50^ We find it, therefore, of interest to compute the *R*_g_ and the end-to-end distance separately for the two domains of FUS for all parameter sets, Fig. 2e,f. The overall trend for both domains was similar to that of the full-length FUS with a few notable exceptions. The ff19SB and aastar parameter sets showed lower than expected average *R*_g_ values (*∼*21 Å and *∼*25 Å respectively) for the fully disordered LC domain, with the highest (*∼*27 Å) value seen for aadisp. For the FUS-RBD, the aastar data show high *R*_g_ average (*∼*35 Å) and the range of *R*_g_ values comparable to that of the CG-KH model. This behavior is consistent with the end-to-end distance data, where the aastar shows higher range of values for RBD as compared to LC and *vice versa* for the aadisp set. While we cannot rule out the effect of insufficient sampling because of the rather short simulation time scale of 5 *µ*s, our analysis suggests that the LC domain is best represented by the aadisp parameter set, while the RBD domain is best represented by the aastar parameter set, both using the TIP4P-D water model.

### 3.2 Intra-molecular interactions and the solvent accessible area

The ability of FUS protein to self-assemble into condensates arises from the amino acid level interactions within the FUS protein and with other FUS proteins. It has been experimentally shown that arginines (R) and tyrosines (Y) in the FUS protein form cation-*π* interactions that drive the phase separation, these residues thus being referred to as stickers.^15^ Other key residues like glycine (G) maintains liquidity, while glutamine (Q) and serine (S) promote hardening of the condensate, thus termed as spacers.^15^

By analyzing our 5 *µ*s trajectories of the full-length FUS protein, we evaluated the efficacy of the nine parameter sets for their ability to replicate these key interactions. Specifically, we determined the 2D self-interaction contact map for all residues of the FUS protein, Fig. 3a, as well as the average number of contacts per simulation time frame along the FUS residues, Fig. 3b. The data for the five most interested parameter sets are shown in Fig. 3, whereas SI Fig. S6 shows data for the other four sets. A characteristic signature of structured interactions is observed at the RNA recognition motif (residues 285 to 371), across all parameter sets, Fig. 3a. Similarly, there are strong structured interactions at the zinc finger domain (residues 421 to 453), except for the tip4pd parameter set, which shows weaker contact interactions, and for the aastar set, where these interactions are even weaker. These observations are even more apparent in the average number of contacts plot (Fig. 3b), where the tip4pd and aastar sets have lower average contacts in the RRM (red background) and ZnF (blue background) domains, as compared to the other parameter sets.

**Figure 3:**
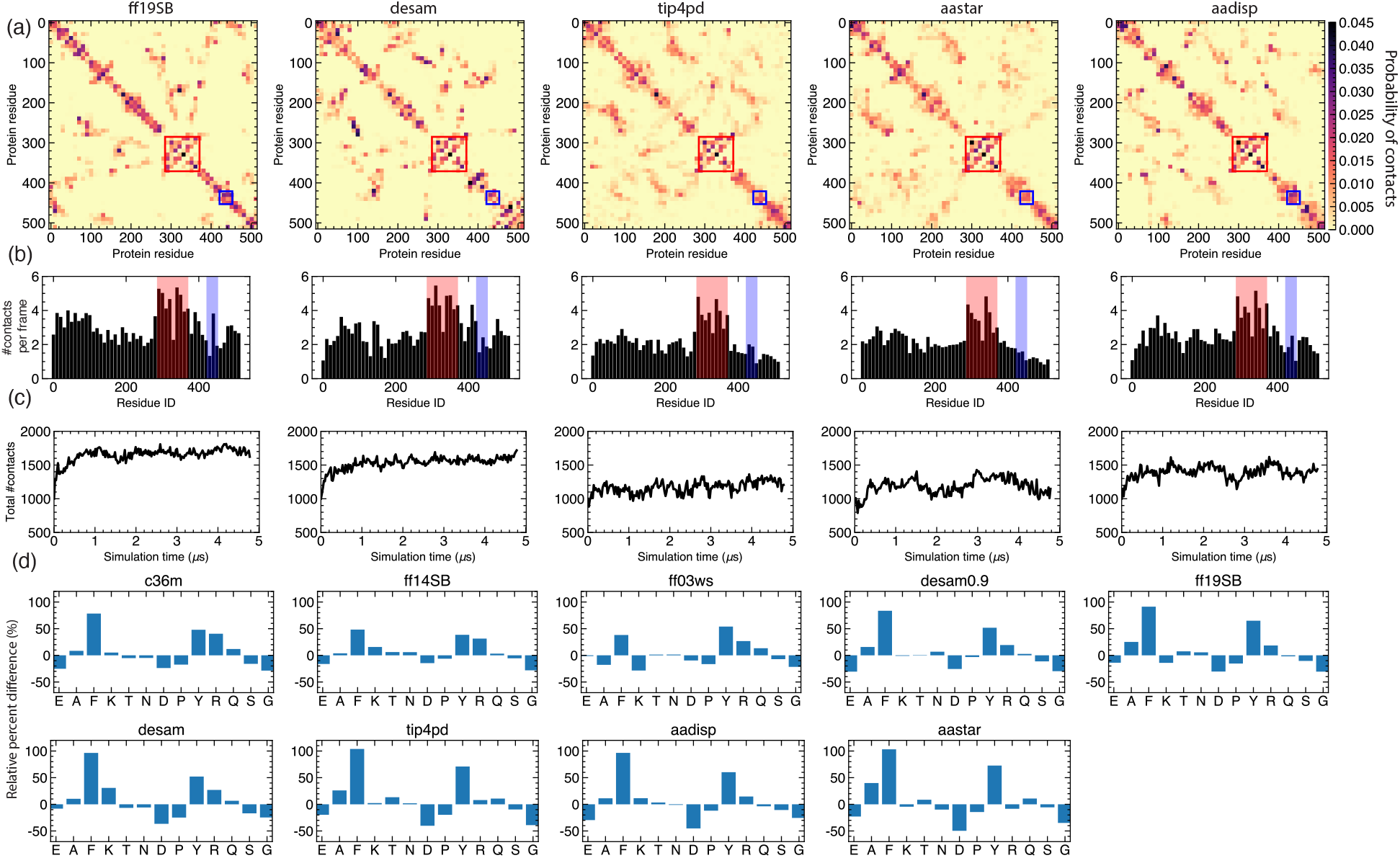
Intramolecular interactions within FUS. (a) Residue-based 2D maps of self contacts extracted from the full-length FUS simulation trajectories. Each data point on the 2D map is block averaged over 10 residues in X and Y axis for clarity and the average contact probability is plotted according to the color bar shown on the right. Two residues are considered to be in contact if any atom of one amino acid is located within 3 Å of any atom of the other residue; nearest three neighbors are excluded from the analysis. The structured regions of FUS are highlighted in red (RRM) and blue (ZnF). (b) Average number of contacts a given protein residue makes with other residues of FUS (black). Each bar value is block averaged over 10 consecutive residues. The structured regions are highlighted in red (RRM) and blue (ZnF). (c) Total number of unique pairwise intramolecular contacts as a function of simulation time. (d) Relative percent difference between the self-contacts involving a particular amino acid and the abundance of that amino acids in the FUS sequence, normalized by the abundance. Data for amino acids of low (*<*2%) abundance are not shown. The amino acids are arranged in the ascending order according to their abundance in the FUS sequence.

The choice of a force field has also a pronounced effect on the off-diagonal elements of the self-interaction map, Fig. 3a. In the case of ff19SB and desam, the off-diagonal interactions are localized to a few regions, indicating the formation of sticky contacts in the disordered domains that persist over the course of the entire simulation trajectory. In contrast, the off-diagonal regions are diffuse in the tip4pd, aastar and aadisp maps, Fig. 3b, indicating transient and less abundant interactions within the disordered domains. The total number of self-contacts, Fig. 3c, reaches a constant value after approximately 1*µ*s and is the largest for the ff19SB and desam sets, followed by aadisp and then by tip4pd and aastar. Overall, a larger average *R*_g_ value correlates with a smaller number of self-contacts, although the dependence is more nuanced, as a comparison of the aadisp and aastar results show.

Through further analysis (SI Fig. S6d), we find the following five residues arginine (R), tyrosine (Y), glutamine (Q), serine (S) and glycine (G), to contribute *∼*60% of all self-interactions. The high abundance of interactions involving arginines and tyrosines is consistent with their expected role of ”stickers” at the intra-protein level being the driving forces for phase separation. The glutamine and serine amino acids have been classified as ”spacers” and their dominant contribution to intra-FUS self-interactions alludes to a contact-based mechanism for condensate hardening. Interestingly, glycine has emerged as the most abundant residue forming the network of self-contacts, previously termed as spacer^15^ owing to its conformational flexibility and being one of the frequent ALS mutant sites.^51^ We note, however, that the frequent occurrence of self-contacts involving the R, Y, Q, S, and G residues in hardly surprising as these residues are also the most abundant in the FUS sequence, Fig. 4c.

**Figure 4:**
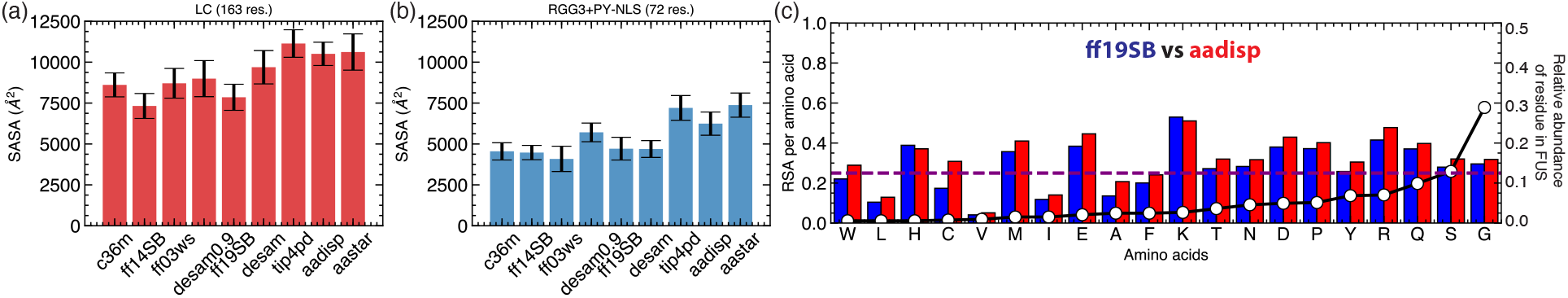
Solvent accessibility of FUS domains. (a) Solvent accessible surface area (SASA) for the disordered FUS-LC domain across the nine parameter sets. SASA was calculated using VMD, ^73^ a 1.4 Å radius probe and averaged over snapshots taken every 24 ns over the entire simulation trajectory. The error bars represent the standard deviation over the snapshots. (b) Same as in panel a but for the disordered FUS tail containing the RGG3 and PY-NLS domains. The error bars represent standard deviations over simulation trajectory. (c) Relative solvent accessibility of individual amino acid types in the ff19SB (blue) and aadisp (red) runs. The dashed purple line shows the threshold (0.25) below which a residue is considered to be buried. The black line with white circles (right axis) shows the relative abundance of each amino acid type in the FUS protein.

To remove the compositional bias from our self-contact analysis, we’ve computed the difference between the fraction of self-contacts involving a particular amino acid type and the relative abundance of that amino acid in the FUS sequence and then normalized the difference by the abundance. The resulting relative difference plots, Fig. 3d, show which amino acids have strong propensity to forming self-contacts (positive values) and which avoid them (negative values), and how the choice of a molecular force field affects such interactions. Thus, we find R and Y residues to clearly show characteristics of a sticker, forming more contact than expected from their abundance. Arginine residues were found to be more stickier when described using the conventional (c36m/ff14SB) parameter sets than in the simulations that involved TIP4P-D water model (tip4pd/aadisp/aastar), which could be partially explained by the increased *R*_g_ values of the latter. Tyrosine residues, however, follow exactly the opposite trend, being the least sticky in the c36m/ff14SB simulations and forming the most contacts in the tip4pd/aadisp/aastar simulations. Interestingly, another aromatic amino acid, phenylalanine (F), is found to be the stickiest residue of all, even for the high *R*_g_ parameter sets (tip4pd/aadisp/aastar), which was previously conjectured on the basis of phenylalanine aromaticity^20^ but was overlooked in FUS because the low abundance of phenylalanine. The number of contacts for spacers glutamine (Q) and serine (S) were within 20% of the value prescribed by their abundance, with glutamines forming more contacts than expected and serines forming fewer of those. The last spacer, glycine (G), was found to form even fewer contacts than prescribed by its abundance, in comparison to serine. The above analysis shows that the high number of self-contacts formed by spacers is not caused by some intrinsic property of the spacer amino acids but rather can be attributed to their high abundance in the FUS sequence. Among spacers (Q, S and G), the propensity to forming contacts was not dramatically affected by the force field choice or found to correlate with the *R*_g_ value, though a change of 20% was not uncommon (as seen for glycines). The observed force field dependence of the contact propensity for other residues (K, E, P, and D) did not follow any previously calculated physical properties.

To characterize the accessibility of the FUS protein to the surrounding solvent, we computed the solvent accessible surface area (SASA) of FUS domains, Fig. 4a,b and of individual residues, SI Fig. S7. Being a reliable metric of intrinsic flexibility and folding propensity, SASA can also serve as an indicator of a protein’s interactions with other biomolecules.^74–76^ The highly disordered LC domain was found to have the lowest SASA values in the simulations carried out using the low *R*_g_ parameter sets, ff14SB and, somewhat surprisingly, desam0.9, whereas the highest SASA values were seen in the tip4pd, aastar and aadisp simulations, Fig. 4a. Pronounced dependence of SASA on parameter sets was also seen in the C-terminal region of FUS, SI Fig. S7. Specifically, the RGG3 and the PY-NLS domains— essential to the nuclear localization of FUS^77^—were observed to be highly exposed and dynamic in the tip4pd and aastar simulations, followed by the aadisp one, with the average SASA value of the three being considerably higher than those of the remaining six, Fig. 4b. Similar trends were observed for the SASA values of other disordered domains and also at the per-residue level, SI Fig. S7.

To elucidate the effect of the force field on solvent accessibility of individual amino acid types, we performed a relative solvent accessibility (RSA) analysis of the aadisp and ff19SB runs, Fig. 4c. RSA is defined as the average SASA per amino acid normalized by the maximum allowed solvent accessibility, which in turn was calculated empirically through analysis of crystal structures.^78^ An amino acid of an RSA value below 0.25 (purple dotted line in Fig. 4c) is considered buried inside the protein. In comparison to the ff19SB run, using aadisp increased the solvent exposure of most residues. The largest RSA values were observed for the R, K, E and D residues, after discarding from the consideration amino acid types of low abundance (*<*2%) in the protein sequence, Fig. 4c (right axis).

Using the aadisp run, we can make some observations regarding the accessibility of the key residue types in FUS. Even though the sticker residues arginines (R) and tyrosines (Y) make a prominent contribution to self-contacts, the residues remain exposed to the solvent allowing for higher order inter-molecular interactions. Other sticker residues, tryptophan (W) and phenylalanine (F), do not play a significant role within intra-FUS interactions because of their low abundance and being mostly buried. Among the spacers that contribute to condensate hardening, i.e., glutamine and serine, glutamine is much more solvent exposed than serine, which is mostly buried. This could mean that glutamine residues contribute to condensate hardening by modulating protein-solvent and inter-protein interactions, whereas serine residues modulate intra-protein interactions. Finally, the spacer glycine residues, which are known to maintain the condensate liquidity and being the most abundant amino acid in the FUS sequence, are observed to be mostly buried inside the protein. This observation is in line with our previous observation of glycine carrying the highest intra-FUS self-contacts, building a network of contacts that maintains the condensate fluidity.

### 3.3 Representation of the structured domains

The C-terminal RNA binding domain of FUS contains two structured regions, the RNA recognition motif (RRM, residues 285-371) and the zinc finger domain (ZnF, residues 421 to 453), flanked by arginine-glycine (RGG) rich disordered motifs and ends with the nuclear localization sequence (NLS) tail, Fig. 5a. The RRM domain has a structure common to many eukaryotic RNA binding proteins including FUS, TDP-43 and the hnRNP family of proteins implicated in RNA processing and gene expression. ^79^ FUS-RRM has the conventional RRM-β_1_α_1_β_2_β_3_α_2_β_4_ fold, consisting of four β-sheets, interspersed with two *α*-helices. The NMR structure of FUS-RRM has been obtained in complex with a stem-loop RNA,^50^ Fig. 5b. The structural motif that facilitates RNA binding is a unique β*^!^*β*^!!^*-hairpin in the α_1_β_2_ loop, Fig. 5b. The FUS-ZnF domain belongs to the RanBP2-type ZnF family of proteins known to bind single-stranded RNA containing GGU motifs and involved in mRNA regulation.^80^ The ZnF domain contains a typical four-cysteine-residue enclosure of a bound zinc ion and two crossed β-hairpins. The solution structure of FUS-ZnF in complex with a UGGUG ssRNA strand was resolved using NMR, Fig. 5b.^50^

**Figure 5:**
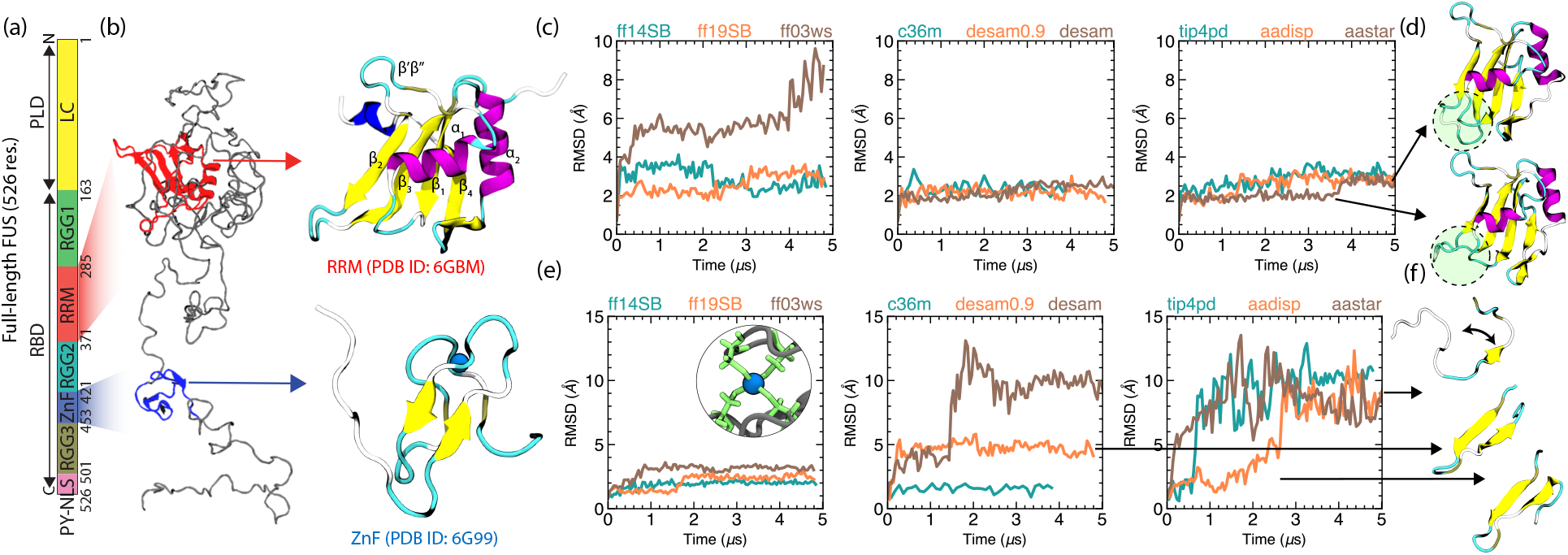
Stability of the structured domains of FUS in the absence of RNA. (a) Domains of a full-length FUS protein annotated by their starting and ending residue numbers using the nomenclature of Yoshizawa *et al.* ^70^ (b) Initial conformation of a FUS protein (grey), where the structured RRM and ZnF domains are highlighted in red and blue, respectively (left). The images on the right provide a zoomed-in view of the NMR structures colored and annotated according to the type of the secondary structure. The zinc ion in the ZnF domain is shown as a blue sphere. The structures were taken from the protein data bank entries 6G99 (ZnF) and 6GBM (RRM). ^50^ (c) Average root mean square deviation (RMSD) of the C*_α_* atoms of the RRM domain (residues 285 to 371) in nine MD simulations of a full-length FUS protein. The RMSD was calculated with respect to the reference pdb structure shown in panel b. (d) The conformations of the RRM’s β_2_β_3_ loop (circle) corresponding to the *∼*2 Å and *∼*3 Å RMSD values, from the simulations indicated by the arrows. (e) Average root mean square deviation (RMSD) of the C*_α_* atoms of the ZnF domain (residues 424 to 443) in nine MD simulations of a full-length FUS protein. The RMSD was calculated with respect to the reference pdb structure shown in panel b. The simulations depicted in the left most figure (ff14SB, ff19SB, ff03ws) were carried out in the presence of explicit covalent bonds between the Zn ion and the sulphur atoms of the four neighboring cysteines, illustrated by the inset image (cysteine in green, zinc in blue). (f) Representative conformations of the ZnF domain (residues 424 to 443, blue) corresponding to a folded structure (*∼*2 Å RMSD, bottom), partially destabilized conformation (*∼*5 Å RMSD, middle) and of a fully unfolded conformation (*∼*10 Å RMSD, top). The highest RMSD states are associated with the opening of the *β*-hairpin, as illustrated by the arrow.

To evaluate the integrity of the two structured domains over the course of our 5 *µ*s equilibration simulations, we computed the root mean square deviation (RMSD) of the domains’ C*_α_* atoms, aligning each domain individually to its respective experimental conformation, Fig. 5c,e. As both the NMR structures and the domain definitions include partially disordered regions, we selected, for our alignment and RMSD analysis procedures, only those residues that are known to be not disordered according to the UniProt Knowledgebase (FUS: P35637). Specifically, residues 285 to 371 were used for the analysis of RRM and residues 425 to 443 for the analysis of ZnF.

Although all our benchmark simulations of the full-length FUS were carried out in the absence of stem-RNA bound to RRM, we observed little to no destabilization of that domain. That is, in eight of out nine simulations, the structure of RRM was maintained with RMSD values below 3 Å, Fig. 5c. The surprising exception was the simulation carried out using the ff03ws set, where the RRM domain was observed to partially unfold, reaching an RMSD of 8 Å. In most simulations, we observed an RMSD shift between 2 to 3 Å, which, on close inspection, was associated with a conformational transition in the β_2_β_3_ loop, as illustrated in Fig. 5d. Interestingly, in the tip4pd, aadisp and aastar simulations, the β*^!^*β*^!!^* hairpin, which interacts with stem-loop RNA only in the FUS-RRM domain, was more stable than the β_2_β_3_ loop, which mediates RNA binding in other RRM containing proteins. ^50^

The removal of ssRNA from the NMR structure of the ZnF domain combined with different methods of enforcing the zinc-cysteine bonds has led to pronounced differences in the simulation outcomes regarding stability of the ZnF domain, Fig. 5e. In the ff14SB and ff19SB simulations, the ZnF domain maintained its integrity, which we attribute to the covalent bonds that we introduced to describe the zinc–cysteine sulphur interactions (see Methods). Despite having such bonds, slight structural destabilization was observed in the ff03ws simulation, where the RMSD value reached 3.5 Å, Fig. 5e. Among the remaining cases, ZnF maintained its integrity during a 2 *µ*s c36m run and partially destabilized during the 5 *µ*s desam0.9 run. The partial destabilization corresponds to a conformational change in 2-3 residues at each end of the ZnF sequence (residues 424 to 443), Fig. 5f. The primary secondary structure element, the β-hairpin, remains folded and maintains its structure throughout the simulation. In the tip4pd and aastar runs, ZnF fully unfolds almost immediately, whereas during the desam and aadisp runs, ZnF remains stable for the first 1.5 and 2.5 *µ*s, respectively. The complete unfolding of ZnF (*∼*10 Å RMSD) in all four cases is associated with the opening of the β-hairpin, which leads to a more extended conformation, Fig. 5f.

We revisit stability of the RRM and ZnF domain in the presence of bound RNA in the next section.

Just like ordered parts of the protein can become disordered, so can initially disordered parts of the protein become transiently or permanently ordered. To investigate this possibility, we used STRIDE^81^ and VMD^82^ to carry out secondary structure analysis on our FUS trajectories for the following five parameter sets: ff19SB, desam, tip4pd, aadisp and aastar. Consistent with the above data, the two major structural elements are found in the RRM and ZnF regions of FUS, which are the only two permanently structured domains, SI Fig. S8a. However, a more detailed view of the fully disordered LC domain in the ff19SB simulation, SI Fig. S8b, revealed formation of small alpha helices of roughly six to eight residues that persisted on the microsecond timescale, with one of them remaining for the entire five microsecond duration of the simulation. The desam simulation showed fewer secondary structure elements, which were localized to a few residue forming bridges and lasting for 1 to 2 microseconds. The secondary structure formation in the aadisp simulation was similar to that of desam, but with a lower abundance of shorter (100 ns) lifetime. Finally, the tip4pd and aastar simulations had the second lowest and the lowest number of secondary structure elements in the LC domain, respectively, which were similar to that of the aadisp simulation but occurring with a lower probability and persistence. Thus, from the five MD trajectories, the ff19SB one was the only simulation where we observed a significant number of spontaneously formed secondary structure elements in the disordered LC domain. The desam and aadisp simulations were characterized by the formation of some transient secondary structures, whereas those were minimal in the tip4pd and aastar simulations.

### 3.4 Binding of RNA to the structured domains

To further characterize the properties of the structured domains, we simulated the RRM and the ZnF domains in the presence of their RNA targets, as captured by their respective NMR structures. In contrast to our previous simulations of the full-length FUS systems, this set of simulations featured only the RRM (residues 276 to 377, PDB:6GBM) or ZnF (residues 414 to 454, PDB:6G99) domain of FUS.

The structure of RRM was resolved by NMR in complex with a 23-nucleotide stem loop RNA, as shown in Fig. 6a. The structure was solvated in a cubic volume of 150 mM KCl solution and, upon minimization and a brief 10 ns restrained equilibration, was simulated for 10 *µ*s in the absence of any restraints using either the ff14SB, tip4pd, aadisp or aastar parameter sets on Anton 2, see Methods and Table 1 for further simulation details. Snapshot in Fig. 6b illustrate the resulting aadisp and aastar trajectories, aligned using protein coordinates as a reference.

**Figure 6:**
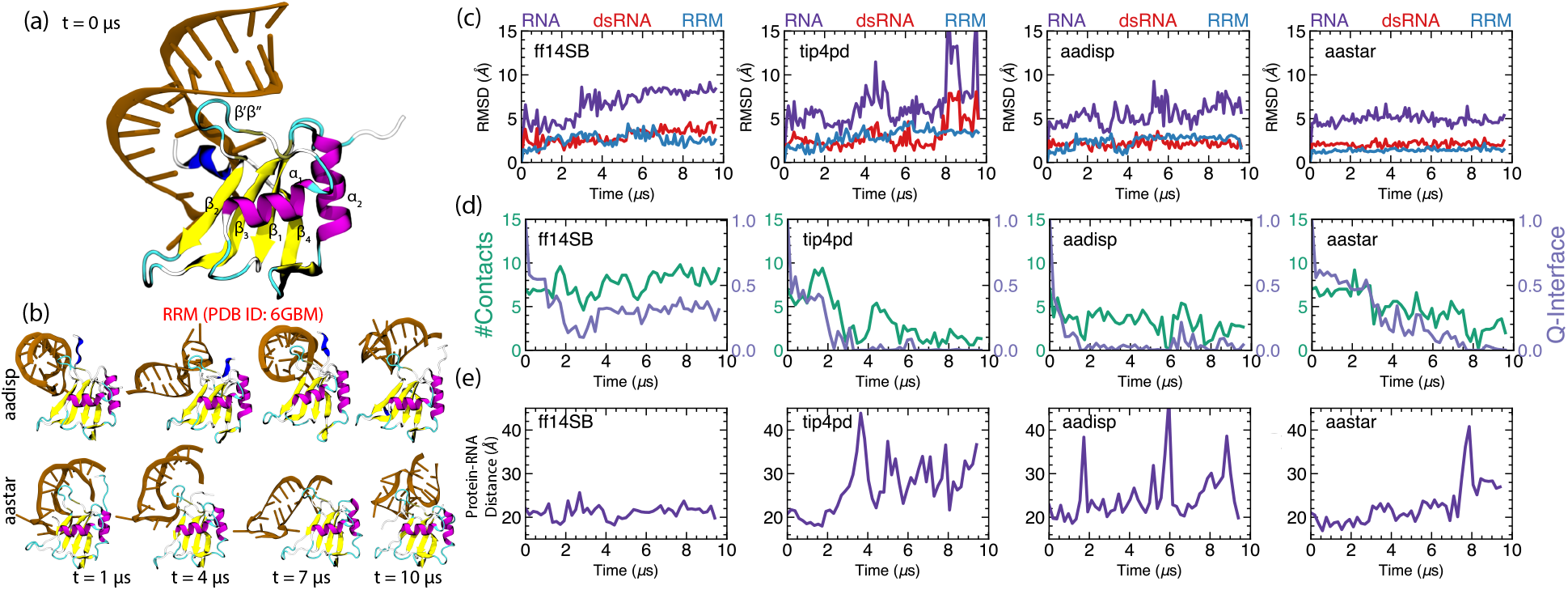
Properties of RRM simulated in complex with its RNA target. (a) NMR structure of FUS-RRM (PDB ID: 6GBM) ^50^ bound to a 23-nucleotide RNA stem loop (brown). The protein is shown using a representation that highlights its secondary structure. (b) Snapshots illustrating two 10 *µ*s simulations of the complex carried out using the aadisp and aastar parameter sets. (c) RMSD of the C*_α_* atoms of residues 285 to 371 of RRM (sky blue), of the non-hydrogen atoms of the sixteen basepair duplex of the RNA (residues 103 to 111 and 118 to 125, red) and of the non-hydrogen atoms of the entire stem-loop RNA (violet) during the ten microsecond simulations carried out using four all-atom force fields (ff14SB, tip4pd, aadisp and aastar). Each RMSD value was calculated with respect to the corresponding coordinates of the reference pdb structure (panel a), upon alignment. (d) The number of protein–RNA contacts (green, left axis) and the q-value of the protein–RNA interface (violet, right axis) as a function of simulation time for the four parameter sets. The number of protein–RNA contacts was computed by counting the number of RNA nucleotides having at least one non-hydrogen atom located within 3 Å of the protein. The q-value of the interface was determined as the fraction of the native contacts remaining relative to that in the PDB structure, as described in Methods. (e) Center-of-mass distance between the RNA and the protein as a function of simulation time for the four parameter sets, in the reference frame of the protein. Periodic boundary conditions were handled by only considering the periodic image of the RNA that was closest to the centered and aligned protein.

In the aastar simulation, the protein remained extremely stable, exhibiting RMSD values below 2 Å, Fig. 6c. Higher (*∼*3 Å) but stable RMSD values were observed for the protein in the aadisp, ff14SB and tip4pd simulations, with the latter exhibiting the largest fluctuations that were less than 5 Å in amplitude and reversible. To evaluate the stability of the stem-loop RNA molecule, we computed the RMSD values separately for its sixteen basepair duplex and for the entire molecule. In all simulations, the RMSD values of the entire RNA molecule were seen to considerably exceed that of its double-stranded part, fluctuating around 5 Å in the aadisp and aastar runs and reaching even higher values in the ff14SB and tip4pd runs, with the latter run featuring a partial denaturing of the stem-loop structure (after 8 *µ*s). The RMSD of the dsRNA, however, remained stable at 2 Å for both aadisp and aastar runs, reaching 4.5 Å for ff14SB and 8 Å for tip4pd, which indicates partial melting of the duplex in the latter run.

Despite both the RRM domain and the dsRNA part of the stem loop showing stable, low-amplitude RMSD values, the entire RNA–protein assembly is found to exhibit a complex, dynamic behavior. Visual inspection reveals the RNA molecule to reconfigure its binding to the protein, Fig. 6b. To quantify this behavior, we computed the total number of contacts between the RNA and the RRM, as well as the fraction of the native contacts (from the NMR structure) remaining at the RNA–protein interface, the so-called Q-interface value, Fig. 6d. The Q-interface value was calculated using the C*_α_* positions of the residues of protein and RNA that are in contact in the PDB structure, using a formula previously described in Winogradoff et. al^68^ and as described in Methods. Only in the case of the ff14SB system, the number of contacts remained constant, with roughly a third of the native contacts remaining throughout the 10 *µ*s simulation. The plot of the center-of-mass RNA–protein distance, Fig. 6e, indicates a stable arrangement of the RNA relative to the protein, suggesting that the complex more-or-less maintains its initial configuration in the ff14SB simulation.

In the remaining three runs, the number of native interface contacts drops to zero with time, with the slowest drop observed for the aastar system, Fig. 6d. At the same time, the total number of protein–RNA contacts drops but does not reach zero. The plot of the protein–RNA distance, Fig. 6e, indicates one unbinding/rebinding event for the aastar run and multiple unbinding/rebinding events for both aadisp and tip4pd runs. It is worthwhile to note that, in both of these systems, the re-association of RNA occur in the same region of RRM— the β*^!^*β*^!!^* hairpin loop (which is bound to RNA in the NMR structure), which, in the aadisp case, interacts with the major groove of dsRNA, just like in the NMR structure. While rapid rebinding by itself is not surprising given the small volume of our simulation system, the repeat targeting of the binding region at the RRM surface is encouraging and suggests that the aadisp model may capture the essence of the RNA-FUS recognition interactions. To note, similar unbinding events were observed in the simulations of the FUS-RNA system by Pokorńa et. al.,^33^ which were done using the ff14SB force field for the protein, the OL3 force field^66^ for the RNA and the OPC water model^41^ along with a set of CUFIX corrections.^37^

Similar simulations were carried out for the ZnF domain which, in its NMR structure, ^50^ is bound to a five-nucleotide ssRNA fragment, Fig. 7a. To note, in our previous simulations of full-length FUS (Fig. 5), we observed unfolding of the ZnF domain for the tip4pd, aadisp and aastar parameter sets. We hypothesized that the primary reason for the unfolding was the inability of the force fields to maintain the coordination of the zinc ion by the sulphur atoms of the four cysteine residues. To eliminate this possibility, the simulations of the ZnF– RNA complex were carried out having the zinc ion and the sulphur (SG) atoms of the four neighboring cysteines (residues 428, 433, 444 and 447) restrained to their NMR coordinates, which maintained the coordination of the zinc ion throughout the simulation.

**Figure 7:**
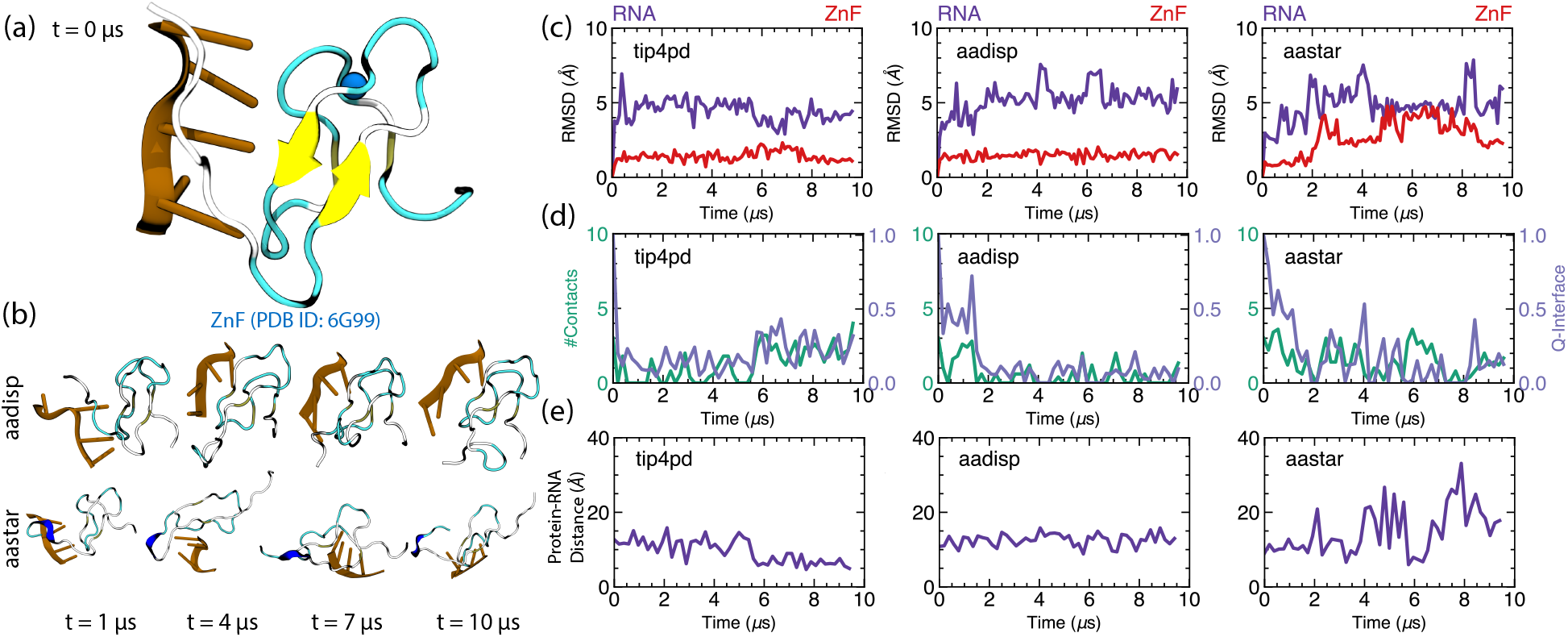
Properties of ZnF simulated in complex with its RNA target. (a) NMR structure of the FUS-ZnF (PDB ID: 6G99^50^) bound to a UGGUG RNA segment (brown), with a bound zinc ion (blue sphere). The protein is shown in a representation highlighting the secondary structure. (b) Snapshots illustrating two 10 *µ*s simulations of the complex carried out using the aadisp and aastar parameter sets. (c) RMSD of the C*_α_* atoms of residues 424 to 443 of ZnF (red), and of the non-hydrogen atoms of the entire UGGUG RNA (violet) during the ten microsecond simulations carried out using three all-atom force fields (tip4pd, aadisp and aastar). Each RMSD value was calculated with respect to the corresponding coordinates of the reference pdb structure (panel a), upon alignment. (d) The number of contacts (green, left axis) and the q-value of the interface (violet, right axis) for the protein-RNA complex as a function of simulation time for the three parameter sets. (e) Center-of-mass distance between the protein and the RNA as a function of simulation time for the three parameter sets, in the reference frame of the protein.

Snapshots in Fig. 7b illustrate the resulting aadisp and aastar trajectories, aligned using as a reference the protein coordinates. The ZnF motif is found to be very stable in the tip4pd and aadisp simulations with an RMSD *<* 2 Å. However, in the aastar run, one of the two beta hairpin loops of ZnF unfolds, Fig. 7b, which increases ZnF RMSD to 5 Å, Fig. 7c. Thus, even in the presence of the restraints that maintained zinc-cysteine bound, we observed partial unfolding of ZnF in the aastar run. To note, among the tip4pd, aadisp, aastar simulations of full-length FUS (Fig. 5), ZnF domain unfolded almost immediately in the aastar simulation.

The RMSD values of the ssRNA fluctuated around 5 Å in all three simulations, which we attribute to it being unstructured. The number of regular and native interface contacts decreased rapidly for the tip4pd set, however half of the native contacts remained in the aadisp and aastar runs for the first 2 *µ*s of the simulation, Fig. 7d. Despite the high RMSD values of the RNA, the protein–RNA association appears to be very stable in the tip4pd and aadisp systems, where the protein–RNA distance remained constant throughout the 10 *µ*s runs, Fig. 7e. In contrast, multiple dissociation and rebinding events were observed during the aastar runs, Fig. 7e, which we attribute to partial unfolding of the ZnF domain.

### 3.5 Conformational dynamics and diffusion

Using our set of nine equilibration trajectories of a full-length FUS protein, we evaluated the effect of the force field on the observed diffusion and structural relaxation. First, we calculated the diffusion coefficient of FUS from its center-of-mass mean squared displacement (MSD). To do that, we unwrapped the protein trajectory to remove the periodic boundary jumps and then calculated the square of the protein displacement over a given time interval, which was then averaged by moving the starting time point of the interval from zero to all possible values, in half interval steps, to avoid data correlation. Examples of typical MSD curves are shown in Fig. 8a. Using the slope of the MSD curves and the Einstein’s diffusion equation, we obtained the diffusion coefficient for all nine parameter sets, Fig. 8b. Not surprisingly, the protein diffusion coefficients reflected the dependence of the water viscosity on the water model,^46^ with the FUS diffusion coefficient for all parameters coming with the TIP3P water model being roughly 2-3 times higher than the ones that come with the TIP4P-D or OPC water models, Fig. 8b. Within each water model, the diffusion coefficients were roughly similar in the order of magnitude. Secondary to that, we observed slight dependence of the FUS diffusion coefficient on the FUS radius of gyration, with more compact conformations (desam, desam0.9) diffusing faster than those with higher *R*_g_ values (tip4pd/aastar/aadisp).

**Figure 8:**
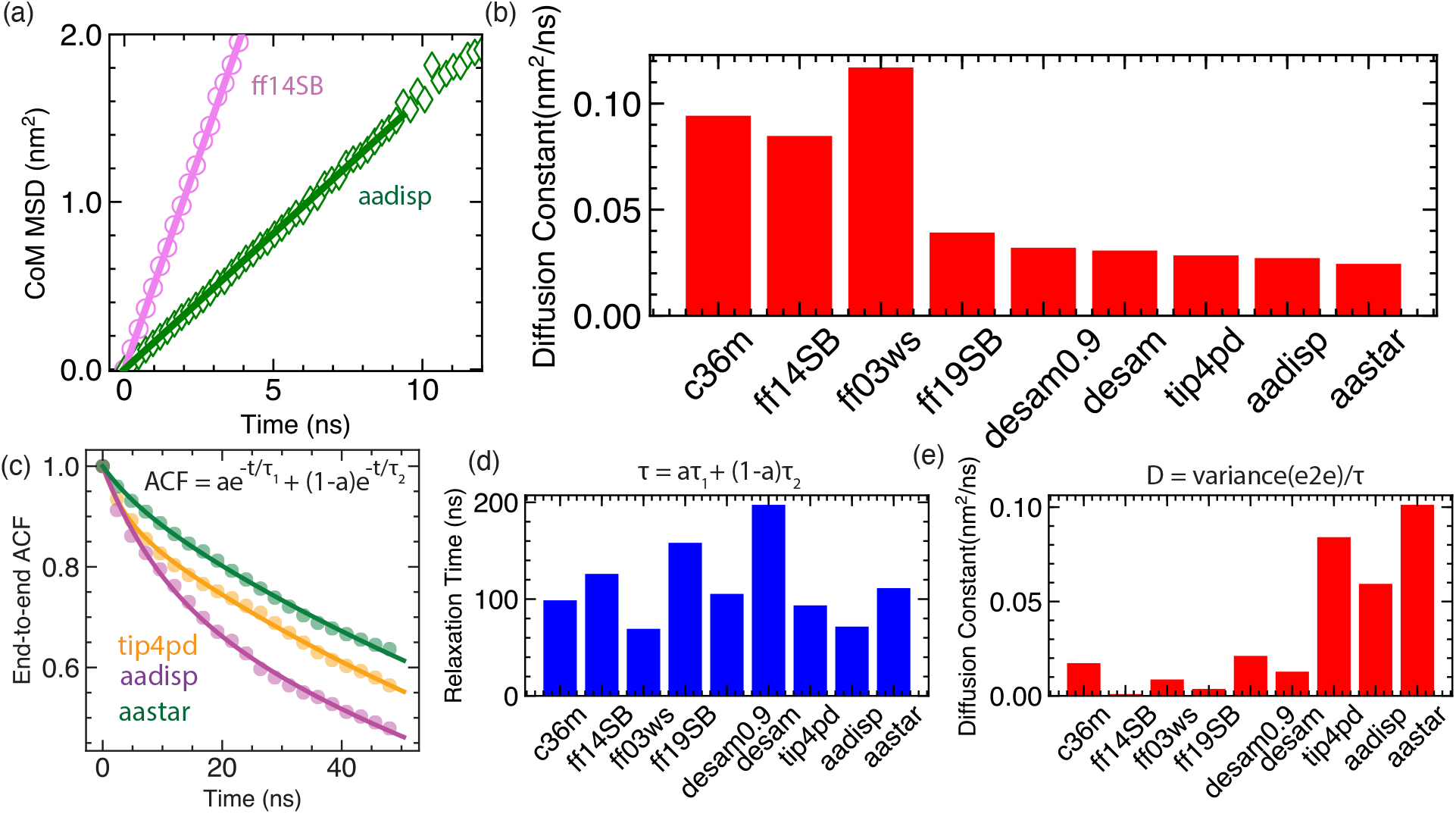
Diffusion and conformational relaxation. (a) Mean square displacement of the FUS protein computed by averaging segments of the 5 *µ*s trajectory for the simulations carried out using the ff14SB (violet) and aadisp (green) parameter sets. (b) The center of mass diffusion constant of the FUS protein calculated from the mean square displacement for nine force-field combinations. Covariance of the FUS end-to-end distance versus the lag time extracted from tip4pd (orange), aadisp (magenta) and aastar (green) trajectories. The first 100 ns of the simulation data were excluded from the analysis. (d-e) End-to-end relaxation time (d) and the diffusion coefficient (e) calculated from the end-to-end distance covariance.

Another approach to obtain the diffusion constant of FUS is to examine the normalized autocorrelation function of its end-to-end distance. Following the method of Benayad et. al.,^83^ we fit the normalized autocorrelation function to a biexponential function as shown in Fig. 8c. The two amplitudes of the exponents were chosen to have the normalized auto-correlation function start at unity at the zero time point. Next, we obtained the effective relaxation time for each parameter set trajectory as an amplitude-weighted average of the two relaxation times, Fig. 8d. Finally, the effective diffusion coefficient was obtained as the ratio of the end-to-end distance variance and the effective relaxation time,^84^ Fig. 8e. Calculated using this method, the diffusion constants and the relaxation times depicts the range and the timescale of the internal motion within the FUS protein, respectively. Interestingly, we find the diffusion constants calculated from the autocorrelation functions to follow an opposite trend to the diffusion constants obtained from the analysis of the mean squared displacement. We hypothesize that the parameter sets that increased FUS *R*_g_, such as tip4pd, aastar and aadisp, also facilitate structural fluctuations within FUS lowering the internal relaxation times. A the same time, proteins of larger *R*_g_exhibit a slower center-of-mass diffusion. Taken together, our analysis illustrates non-trivial effects that a choice of a parameter set could have on the diffusive motion of a protein, and the need to distinguish global center-of-mass displacement from internal fluctuations in the diffusion constant analysis.

### 3.6 Implementation of a99SB-disp force-field in NAMD

Thus far, we have used a custom supercomputer system, D.E. Shaw Research Anton 2, to carry out benchmark simulations of a biological condensate system. At the time of writing, the two best performing force field—asdisp and aastar—have not been widely available outside Anton 2 machines. Furthermore, the very hardware architecture of Anton 2 limits the size of an all-atom system that can be run to approximately 4 million atoms with a recommended upper bound of 700,000 atoms,^52^ which caps the size of a biological condensate system that can be investigated using the explicit-solvent MD method to tens of individual condensate molecules.

To facilitate broader use of the best performing force field and enable large-scale simulations of biological condensates we developed a workflow, Fig. 9a, for using the aadisp and aastar parameter sets within a popular MD engine NAMD2. ^53^ Starting from a collection of PDB files containing atomic coordinates of all non-hydrogen atoms of individual molecular species (protein, water, ions and, optionally, RNA), the *tleap* AmberTool is used to generate library files for each molecular component of the system. A custom *tleap* script is then used to combine those four files into an AMBER-format parameter/topology file (prmtop) and a coordinate file (rst7). The resulting prmtop file is modified to add additional dihedral angle potentials for the protein backbone using the *ParmEd* python module. The modified prmtop and rst7 files are then used directly as input to NAMD2. As an illustration of our workflow, we provide all the files needed to create one system containing one FUS protein in electrolyte solution and another system containing six FUS proteins, three ssRNA fragments, TIP4P-D-1.6 water and ions, as well as the step-by-step instructions for creating such files (Supporting Notes 1 and 2).

**Figure 9:**
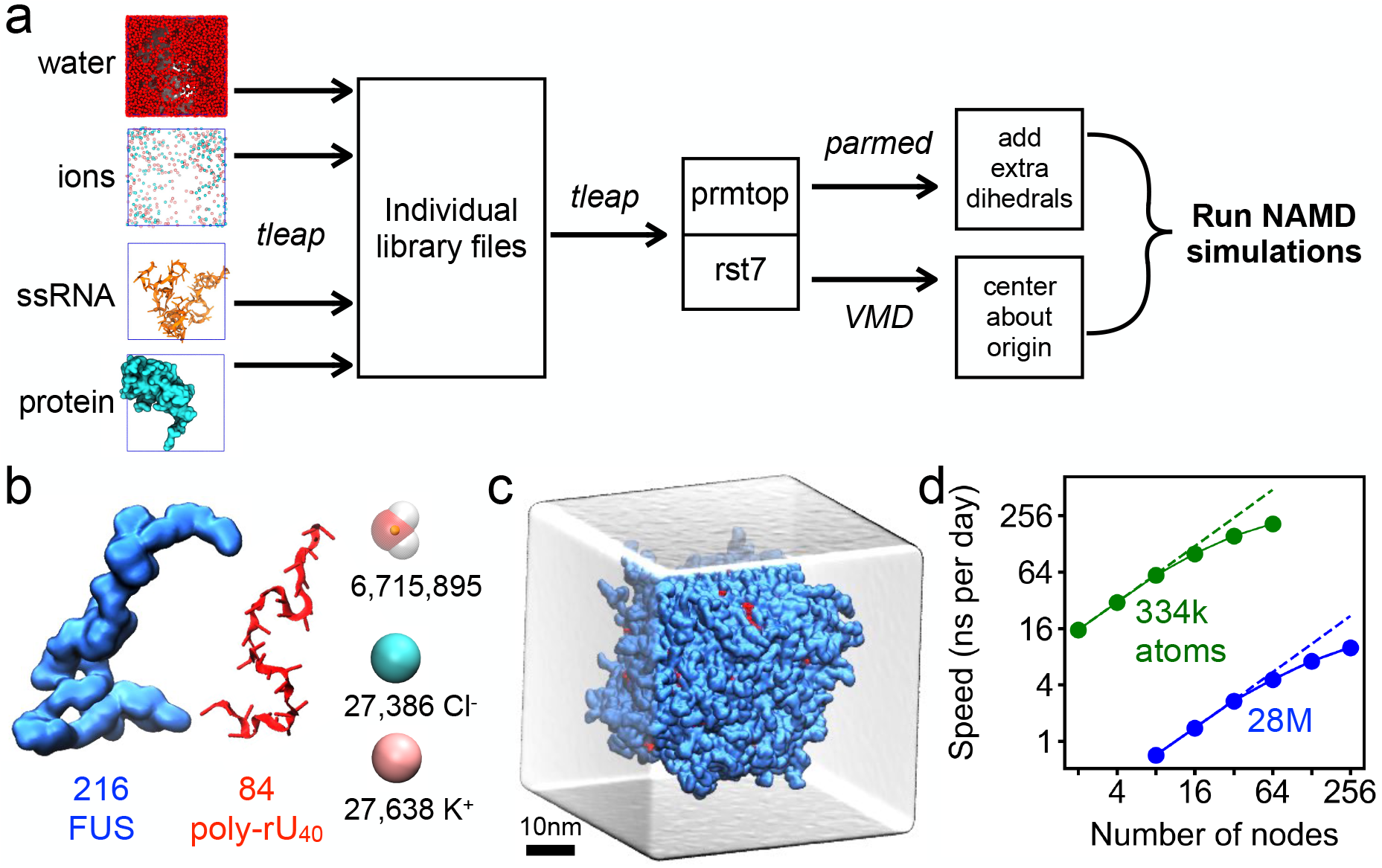
Using a99SB-disp in NAMD. (a) Workflow for preparing a99SB-disp input files for MD simulations using NAMD. Starting from the initial coordinates of the system’s components (PDBs), AMBER-based parameter/topology and coordinate files are generated to be used directly for NAMD runs. (b) Components of the condensate system: 216 FUS proteins and 84 poly-(rU)_40_ RNA strands, 27,638 K^+^ and 27,386 Cl*^−^* ions and 6,715,895 TIP4P-D-1.6 water molecules. (c) All-atom model of a biological condensate, where FUS proteins are shown in light blue, ssRNA in red and the volume occupied by 200 mM aqueous solution of KCl as a semi-transparent surface. The system contains 28,550,760 atoms. (d) Performance of NAMD 2.14 on TACC Frontera a function of nodes used for the simulation of the single FUS system (334,215 atoms, green) and of the RNA/FUS condensate droplet (28,550,760 atoms, blue). The simulations employed a99SB-disp ^48^ for protein, modified AMBER ff14^67^ for ssRNA, CHARMM22^27^ for ions, and the TIP4P-D-1.6^48^ four-point water model. Each node on Frontera contains 56 Intel Xeon Platinum 8280 cores, interconnected by a Mellanox Infiniband HDR, further details in Ref. 85.

We tested our NAMD implementation of the amber99SB-disp force field using the following two biological condensate systems. The first system contained one full-length FUS protein submerged in electrolyte solution, the same 334,215-atom system used for our Anton 2 benchmark runs, Fig. 1a. The second system was a realistic model of a biological condensate formed by 216 FUS molecules, 84 poly(rU)_40_ RNA strands and 200 mM KCl electrolyte, Fig. 9b, a system of over 28 million atoms. The initial all-atom model of the condensate was built by back-mapping an equivalent system’s configuration resulting from a coarse-grained MD simulations of the condensate.^86^ Using NAMD2 on the Frontera super-computer at the Texas Advanced Computing Center, we achieved over 150 ns/day for the system containing one FUS protein, running on 64 nodes, Fig. 9d. For the larger condensate system, close to linear scaling is observed when using up to 128 nodes, Fig. 9d. Note that the 28 million atom system is near the upper limit of the system size that NAMD2 can run without using the memory-optimized input file formats.^87^ Not using a memory-optimized version of NAMD has also affected the overall performance of the 28-million-atom simulation.

### 3.7 Validation of NAMD implementation

We validated our implementation of the aadisp force field in NAMD by simulating the one-FUS system for 5 *µ*s on Frontera (the run referred to aadispNAMD), and comparing the resulting properties of the FUS molecule to those observed in the 5 *µ*s run of the same system using the same force field on Anton 2 (the run referred to as aadispAnton). For comparison, we used NAMD to simulate the same system with the CHARMM36m force field (denoted as C36m).

All three simulations started having the FUS protein in the same extended configuration, Fig. 10a; Fig. 1a shows the same configuration along with the surrounding solvent. By visualizing the final conformation of FUS after a 5 *µ*s run, Fig. 10b, we find that both aadispNAMD and aadispAnton runs produce a more compact configurations than the initial one and that the C36m run produces an even more tightly packed configuration. The plot of protein’s *R_g_*as a function of time, Fig. 10c, indicates that a partial collapse of the FUS conformation occurs within the first microsecond of each simulation, and that aadispNAMD and aadispAnton runs produce similar *R*_g_ values whereas the *R*_g_ values from the C36m run are much smaller. The protein’s end-to-end distance, Fig. 10d, samples a wide range of values in both aadispNAMD and aadispAnton runs, but drops and stays low in the C36m run, similar to the behavior observed in the CHARMM36m simulation of FUS on Anton 2, SI Fig. S1 and S4.

**Figure 10:**
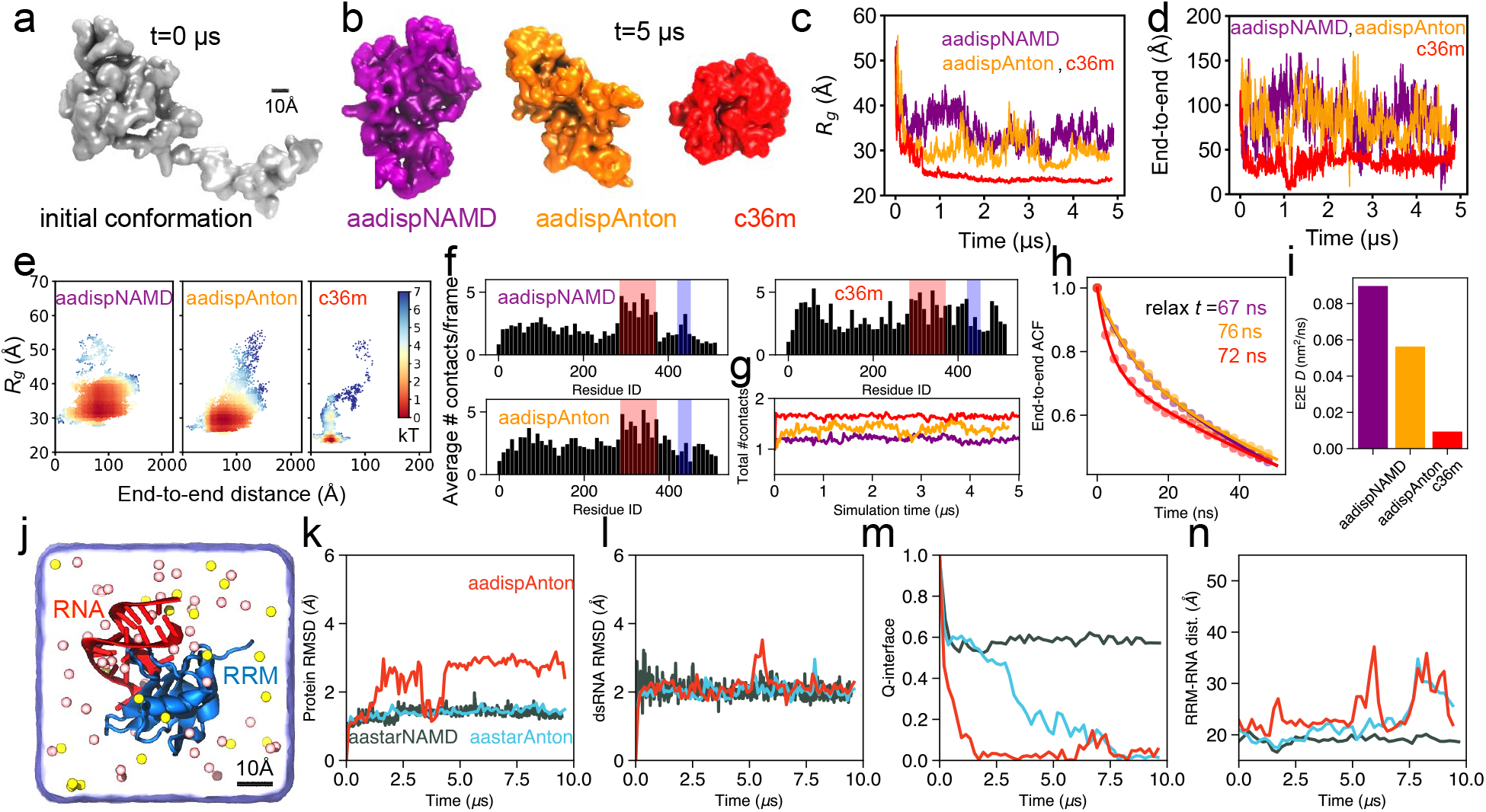
Validation of aadisp implementation in NAMD. (a) Starting configuration of FUS. (b) Configuration of FUS after 5 *µ*s of MD simulation at 310 K. The a99SB-disp force field simulations performed using NAMD and Anton 2 are denoted as aadispNAMD (purple) and aadispAnton (orange); c36m (red) stands for CHARMM36m. (c,d) Radius of gyration (panel c) and end-to-end distance (panel d) of the FUS protein as a function of simulation time. (e) Free energy maps of the FUS protein’s conformation as a function of its radius of gyration and end-to-end distance. (f) Average number of self-contacts per frame by FUS residue, calculated in the same manner as Fig. 3b. (g) Total number of self-contacts within FUS as a function of simulation time. (h) End-to-end auto-correlation function *versus* lag time (excluding the first microsecond of simulation). (i) End-to-end diffusion coefficient, calculated as *σ*^2^*/τ*, the variance divided by the relaxation time. (j) Solvated system containing the NMR structure^50^ of FUS-RRM (blue) bound to RNA (red). (k) RMSD of the C*_α_* atoms of residues 285 to 371 of RRM for the aastar (aastarNAMD, green) simulation using NAMD. The same values from the aadisp (aadispAnton, red) and aastar (aastarAnton, blue) Anton simulations are reproduced from Fig. 6c. (l) RMSD of the non-hydrogen atoms of the sixteen basepair duplex of the RNA (residues 103 to 111 and 118 to 125) with respect to the pdb structure. (m) Q-value of the protein–RNA interface. (n) Center-of-mass distance between the RNA and the protein, in the reference frame of the protein. The color scheme for panels l–n match that of panel k. The Anton results are reproduced from Fig. 6.

Consistent with the analysis of our Anton 2 simulations, the free-energy landscape explored by the FUS protein in the aadispNAMD run occupies a relatively broad and shallow basin, when mapped using the *R_g_* and end-to-end distance as the map’s coordinates, Fig. 10e. In contrast, the C36m map exhibits a compact free energy well, spanning a relatively small range in both *R_g_* and end-to-end distance values within 2 *k_B_T* of the global minimum. The per-residue plot of the average number of self-contacts, Fig. 10f, show quantitatively similar patterns for aadispNAMD and aadispAnton runs, whereas the data from the c36m run exhibit a pronounced increase in the number contacts in the RNA recognition motifs and in other domains of FUS. Indeed, the plot of the total number of self-contacts as a function of simulation time, Fig. 10g, shows a larger number of such contacts forming in the C36m simulations than in either aadispNAMD or aadispAnton runs, though the last two also exhibit quantitative differences among themself. We attribute the latter differences, as well as those seen in the free-energy maps, Fig. 10e, to intrinsic differences in the paths of the simulation trajectories, Fig. 10c, and insufficient sampling of the protein configurations afforded by the 5 *µ*s simulation time. Finally, to compare the kinetics of FUS in Anton and NAMD simulations, we calculated the autocorrelation function of the end-to-end distance, Fig. 10h, which followed a relatively similar decay versus lag time for all three force fields. Because the aadispNAMD and the aadispAnton end-to-end distance varies more in time compared to C36m, the aadisp force fields have a significantly greater end-to-end diffusion coefficient than C36m, Fig. 10i.

Using the systems described in Fig. 5, two validation simulations were performed to test the description of RNA–FUS interactions within NAMD. We simulated the RRM-RNA system for up to 10 *µ*s using the the aastar (the run referred to as aastarNAMD) parameter set and compared the resulting properties with the Anton simulations. The simulation started from the same conformation as in the protein data bank and the corresponding Anton simulations, Fig. 10j. Comparing the RMSD of the structured protein, the aastarNAMD run shows a highly stable protein with RMSD values below 2 Å, just like in the Anton trajectories, Fig. 10k. The stem loop RNA was exceptionally stable, with the RMSD of the double stranded region remaining at 2 Å, Fig. 10l. By plotting the Q-interface value versus time, Fig. 10m, we find the aastarNAMD run to maintain the native contacts, very similar to the behavior seen in the aastarAnton run, reaching a stable Q-value of 0.6 even after 9 *µ*s of simulation. The protein-RNA distance, Fig. 10n, in the aastarNAMD run was stable, exhibiting less fluctuation and no detachments similar to that observed in the corresponding Anton run over the same timeframe.

Taken together, our test simulations validate our implementation of the aadisp and aastar force fields in NAMD, opening their use to a broader research community.

## 4 DISCUSSION AND CONCLUSION

In this work, we have evaluated performance of nine contemporary force fields (Table 1) for all-atom MD simulations of biological condensates. Using the Anton 2 machine, we performed nine 5 *µ*s benchmark simulations of a 526-residue, full-length protein FUS that contains both structured and disordered regions. Side-by-side comparison of the protein’s radius of gyration, Fig. 1e, showed that conventional protein force fields, AMBER ff14SB and CHARMM36m, produce densely packed globular conformations that lack internal dynamics, Fig. 2a and Fig. 8e. Replacing the conventional three-point water model (TIP3P) with either OPC (ff19SB) or TIP4P(ff03ws) four-point water models slightly expanded the ensemble of conformation, which nevertheless remained considerably smaller than suggested by experiment, Fig. 1f. The introduction of the TIP4P-D water model produced a modest (desam0.9 and desam) to considerable (aastar) increase in the conformational dynamics of the protein, even when the TIP4P-D model was combined with the older ff14SB protein force field (tip4pd). A considerable increase of the *R*_g_ values and enhanced conformational dynamics was also seen for a variant of the TIP4P model, TIP4P-D-1.6 (aadisp). These results indicate that the global properties of the condensate molecule and its internal dynamics are predominantly determined by the choice of the water model, although the effect of the protein force field cannot be discounted.^45^ We also note that none of the all-atom force fields afforded sampling of the conformational ensemble captured by the DLS measurement, implying that the current force fields still leave some room for improvement. In contrast, the ensemble of conformations explored by a coarse-grained model^56^ appears to match the range of conformations captured by the DLS measurements.

The choice of the molecular force field was found to have an expected effect on the number of internal contacts within a FUS molecule, Fig. 3, and the solvent accessible surface area, Fig. 4, both being anticorrelated with the protein radius of gyration. Despite these global quantitative differences, the pattern of local interactions has shown some robust, force field-independent features, including classification of the protein residues to stickers and spacers categories. Analysis of the simulation trajectory obtained using the best performing force field (a99SB-disp) detailed the role of individual amino acids in modulating the physical properties of the condensate.

The force field choice was also found to have contrasting effects on the global diffusion and internal dynamics of a FUS protein. The parameter sets that employed a TIP3P water model were found to exhibit faster Brownian motion relative to the solvent than in the simulations that employed a four-point water model. This observation is fully explained by a factor 3 difference in the water viscosity and the smaller effective hydrodynamic radius of FUS seen in the TIP3P simulations. At the same time, a FUS protein in a TIP3P simulation showed considerably slower internal dynamics, resulting in much lower effective diffusion constants when the latter is estimated from the covariance of the end-to-end distance, Fig. 8e. Among the parameter sets employing a four-point water model, the fastest effective diffusion was observed for the tip4pd, aadisp and aastar parameter sets, offering the most extensive sampling of the protein conformational space.

Whereas high *R*_g_ values and fast internal dynamics are both highly desirable features of an all-atom MD simulation of a biological condensate molecule, those properties alone are not sufficient to guarantee an accurate simulation as biological condensate molecules frequently feature folded regions and exhibit specific iterations with RNA. Thus, we analyzed the ability of the nine force fields to preserve the structure of the folded regions of FUS, Fig. 5, and, separately, carried out several 10 *µ*s simulations of the structured domains bound to their RNA targets, Fig. 6 and Fig. 7. In our nine simulations of the full length FUS, the conventional parameter sets unsurprisingly performed well at maintaining integrity of the structured regions, Fig. 5. All three force fields that produced the highest *R*_g_ values (aastar, aadisp and tip4pd) also maintained the structure of the RNA recognition motif, showing a conformational change in a beta loop essential to nucleic acid binding. The simulations of the zinc finder domain, however, showed immediate (aastar and tip4pd) and delayed (aadisp) unfolding of the domain as those simulations were performed without explicit restrains to maintain the structure of the zinc-cysteine binding pocket.

In our 10-*µ*s simulations of the RRM domain bound to its target RNA, Fig. 6, the structure of the protein was mostly stable in the high *R*_g_ parameter set simulations, exceptionally so in the aastar simulation. Interestingly, for the aadisp set, there were multiple protein-RNA dissociation events, but rebinding of RNA took place at the experimentally observed binding site, showing high specificity. Thus, the aastar and aadisp parameterizations provide qualitatively different description of RNA–protein interactions (stable for aastar and dynamic for aadisp), and would benefit from direct experimental validation. The simulations of the zinc finger domain bound to ssRNA target were performed using explicit restraints on the zinc ion coordination site. In the tip4pd and aadisp simulations, the domain maintained its structure for the full duration (10 *µ*s) of the run, Fig. 7. However, in the aastar simulation, the domain completely unfolded despite the zinc ion restrains, highlighting potential problems in using this parameter set for simulations of partially structured proteins.

Based on the above discussion, we consider both aadisp and aastar to provide considerable advantage over other choices. Interestingly, the combination of an older force field (ff14SB) with an updated water model (TIP4P-D) can be a viable alternative if aadisp and/or aastar parameterization is not available for the system of interest. We note, however, that the structure of RRM became unstable when the ff14SB/TIP4P-D set for the protein was combined with the AMBER OL3 set of the RNA. The outcome of the aadisp and aastar simulations can differ, however, regarding stability of the structured domain (less stable for aastar) and RNA–protein interactions (more stable binding for aastar). As these two force fields are not widely available outside the Anton 2 machine, we have meticulously ported them for use with the NAMD engine. Our NAMD implementation enables simulations of large, multimillion-atom biomolecular condensate systems as well as the application of enhanced sampling methods.

## Supporting information

Files needed to build FUS in electrolyte

Files needed to build a small RNA/FUS condensate

Archive of leap files

## Acknowledgement

This work was supported by the Center for the Physics of Living Cells through the National Science Foundation grant PHY-1430124, and by the National Institutes of Health through grants R01-GM137015 and R21-HG011741 (A.A.) and RF-1AG071326 and RF1-NS113636 (Y.G. and S.M.). Supercomputer time was provided by Leadership Resource Allocation MCB20012 on Frontera at the Texas Advanced Computing Center, and Anton 2 allocation MCB100016P. Frontera is made possible by National Science Foundation award OAC-1818253. Anton 2 computer time was provided by the Pittsburgh Supercomputing Center through grant R01-GM116961 from the National Institutes of Health. The Anton 2 machine was made available by D. E. Shaw Research.

## Supporting Information Available

The Supporting Information contains step-by-step instructions for building a99SB-disp systems for NAMD simulations containing either a single FUS protein or a small volume of an FUS–RNA condensate; description of procedures used for validation of the NAMD implementation of the a99SB-disp force field, results of additional DLS experiments; additional plots of *R*_g_, end-to-end distance, protein contacts, SASA data and secondary structure elements in FUS. This information is available free of charge via the Internet at http://pubs.acs.org.

## Graphical TOC Entry

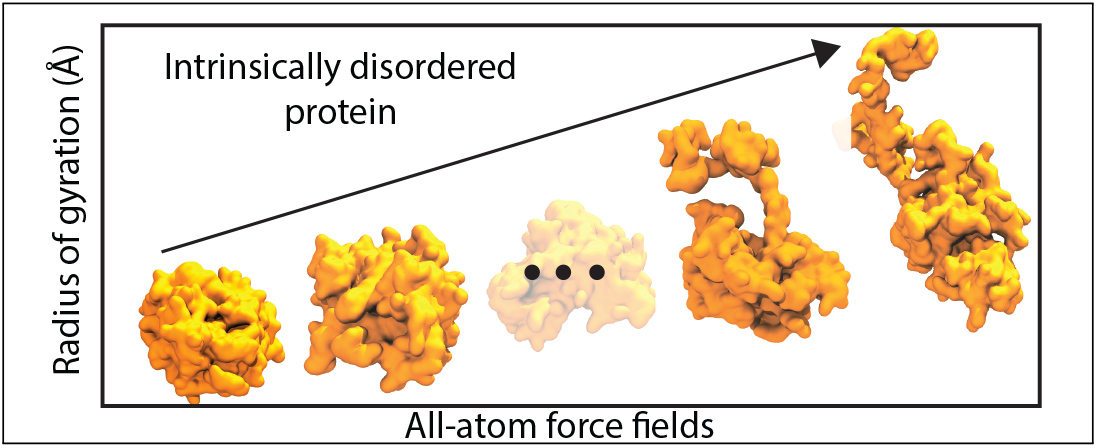

