## Supplementary Information for "Benchmarking Molecular Dynamics Force Fields for All-Atom Simulations of Biological Condensates"

### Supporting Information Note 1: Building an a99SB-disp system containing one FUS protein in solution for NAMD runs

We consider our system of interest containing one FUS protein (protein, water, ions). File names, some content from files, and inputs into the command line will be written in typewriter font. The starting point was three PDBs without hydrogens, taken from the Anton system: (1) `noh_protein.pdb`, (2) `noh_water.pdb`, (3) `ions.pdb`, We implemented the following force fields: a99SB-disp[1] for proteins, CHARMM22[2] for ions and the TIP4P-D-1.6[1] four-point water model. We used AmberTools,[3] VMD,[4] and python scripts to build a parameter-topology (prmtop) and coordinate file (rst7) which can be used directly as input for a NAMD simulation. AmberTools and VMD must be installed, and an Amber environment must be set to proceed. We provide the process as Steps 1–7 below. All the required files are in the `building_1FUS.zip` file, with the force field files in the `amber_leap_files.zip` file.

**(Step 1) Protein**, we start from "`noh_protein.pdb`", which contains one FUS molecule. An AmberTool tleap script was run (`>> tleap -f shaw_fus_leaprc.sh`) to convert those PDBs into Amber-format library files "`shaw_fus.lib`".

**(Step 2) Ions**, start from "`ions.pdb`", which contains zinc, potassium and chloride ions, respectively named "`ZN`", "`K+`" and "`Cl-`". The tleap script was run
(`>> tleap -f ions_leaprc.sh`), to generate the library file "`ions.lib`".

**(Step 3) Water**, start from "`noh_water.pdb`", which contained a single oxygen atom for each water molecule. Each residue was named "`WAT`" and each oxygen atom "`O`". A python script was run (`>> python water_readpdb.py`), to renumber the residues and reformat the PDB, creating "`renum_O_only_water.pdb`". The tleap script was then run

(>> tleap -f shaw_water.sh), to generate the library file "water.lib". Note, we edited the file that typically defines TIP4P-EW, "leaprc.water.tip4pew", to define the four-point water model TIP4P-D-1.6.

**(Step 4) Combine the system together.** We then had three library files: (1) shaw_fus.lib, (2) water.lib, (3) ions.lib. They were combined with the following tleap script (>> tleap -f combine_protein_rna_water_ions.sh),
which generated "protein_rna_water_ions.{prmtop,rst7}".

**(Step 5) Add extra backbone dihedrals to the protein.** We used the ParmEd package in python to complete multi-component dihedrals defined in the a99SB-disp protein force field. The python script (>> python add8.py) was run to generate "add8_protein_rna_water_ions.prmtop".
The script for this step has been written specifically for a single protein (FUS) system, and we ask the user to be careful while using it for a different system and if necessary, update the part commented as "system specific information".

**(Step 6) Center the system, write PDBs for restraints (optional).** Before running, we completed two optional steps in VMD: (1) center the system about the origin in, and (2) write PDB files to restrain non-hydrogen protein and RNA atoms or to restrain C-alpha and C1-prime atoms.

**(Step 7) Run the system with NAMD.** We then ran a series of NAMD simulations, starting from the Amber-format files, "centered_add8_protein_rna_water_ions.{prmtop,rst7}". This included minimization, restrained equilibration, and production simulation runs.

### Supporting Information Note 2: Building a nucleoprotein system for NAMD runs with a99SB-disp force field for proteins and modified Amber ff14 force field for RNA

As an example, here we consider a system containing protein, single-stranded RNA, water, and ions. File names, some content from files, and inputs into the command line will be written in typewriter font. The starting point was four PDBs without hydrogens: (1) `noh_protein.with_RNA.pdb`, (2) `noh_nucleic.with_RNA.pdb`, (3) `ions.with_RNA.pdb`, (4) `noh_water.with_RNA.pdb`. We implemented the following force fields: a99SB-disp[1] for proteins, modified Amber ff14[5] for single-stranded RNA, CHARMM22[2] for ions and the TIP4P-D-1.6[1] four-point water model. We used AmberTools,[3] VMD,[4] and python scripts to build a parameter-topology (prmtop) and coordinate file (rst7) which can be used directly as input for a NAMD simulation. AmberTools and VMD must be installed, and an Amber environment must be set to proceed. We provide the process as Steps 1–8 below. All the required files are in the `building_6FUS_3RNA.zip` file, with the force field files in the `amber_leap_files.zip` file.

**(Step 1) Protein**, we start from "`noh_protein.with_RNA.pdb`", which contains six FUS molecules. A tcl script was run in VMD
(`>> vmd -dispdev text -e write_separate_protein_files.tcl`),
to write six separate files "`fus.{1..6}.pdb`". An AmberTool tleap script was then run (`>> tleap -f shaw_fus_x6_leaprc.sh`) to convert those PDBs into Amber-format library files "`shaw_fus{1..6}.lib`". The following tleap command
(`>> tleap -f shaw_combine_6fus_leaprc.sh`) was then used to combine those six library files into one, "`shaw_protein.lib`".

**(Step 2) RNA**, we start from "`noh_nucleic.with_RNA.pdb`", which contains three segments of ssRNA. A tcl script was run in VMD

(>> vmd -dispdev text -e write_separate_nucleic_files.tcl),

to write three separate files "rna{1,2,3}.pdb". The residues in each separate PDB were renumbered to start from one, "renum_rna{1,2,3}_R.pdb". Each of these ssRNA's were poly-rU$_{50}$, non-terminal residues with the name "U", the 5-prime as "5U" and 3-prime as "3U". A tleap command

(>> tleap -f shaw_rna_x3_leaprc.sh) built three library files "shaw_rna{1,2,3}.lib". Then the following command (>> tleap -f combine_3rna_leaprc.sh) combined those three library files into one, "shaw_rna_x3.lib".

**(Step 3) Ions**, start from "ions.with_RNA.pdb", which contains zinc, potassium and chloride ions, respectively named "ZN", "K+" and "Cl-". The tleap script was run

(>> tleap -f shaw_ions_leaprc.sh), to generate the library file "shaw_ions.lib".

**(Step 4) Water**, start from "noh_water.with_RNA.pdb", which contained a single oxygen atom for each water molecule. Each residue was named "WAT" and each oxygen atom "O". A python script was run (>> python water_readpdb.py), to renumber the residues and reformat the PDB, creating "renum_noh_water.with_RNA.pdb". The tleap script was then run

(>> tleap -f shaw_water_redone_leaprc.sh), to generate the library file "shaw_water.lib". Note, we edited the file that typically defines TIP4P-EW, "leaprc.water.tip4pew", to define the four-point water model TIP4P-D-1.6.

**(Step 5) Combine the system together.** We then had four library files: (1) shaw_protein.lib, (2) shaw_rna_x3.lib, (3) shaw_water.lib, (4) shaw_ions.lib. They were combined with the following tleap script

(>> tleap -f shaw_combine_protein_rna_water_ions.sh),

which generated "shaw_protein_rna_water_ions.{prmtop,rst7}".

**(Step 6) Add extra backbone dihedrals to the protein.** We used the ParmEd package in python to complete multi-component dihedrals defined in the a99SB-disp protein force field. The python script (>> python add8.py) was run to generate "`add8_shaw_protein_rna_water_ions.prmtop`".

The script for this step has been written specifically for our six protein (FUS) system, and we ask the user to be careful while using it for a different system and if necessary, update the part commented as "system specific information".

**(Step 7) Center the system, write PDBs for restraints (optional).** Before running, we completed two optional steps in VMD: (1) center the system about the origin in, and (2) write PDB files to restrain non-hydrogen protein and RNA atoms or to restrain C-alpha and C1-prime atoms.

**(Step 8) Run the system with NAMD.** We then ran a series of NAMD simulations, starting from the Amber-format files, "`centered_add8_shaw_protein_rna_water_ions.{prmtop,rst7}`". This included minimization, restrained equilibration, and production simulation runs.

### Supporting Information Note 3: Verification of the a99SB-disp force field implementation.

We verified our implementation of the a99SB-disp force-field by matching the a99SB-disp implementation on Anton 2, which we considered to be our gold standard for comparison. The following steps can be used in general for comparing force field implementations with their corresponding Anton implementations. We ask the reader to be careful with respect to the versions of the Anton conversion scripts as they are subject to change with regular updates on Anton. We started from AMBER format files: parameter/topology file "`start.prmtop`" and coordinate file "`no-v-start.rst7`". We provide the process below as Steps V1–V3.

**(Step V1) Add velocities to the coordinate file.** We ran a short NAMD simulation to obtain velocities, "`file.vel`". We then loaded the `prmtop` file into VMD, as well as the the NAMD binary file "`file.vel`". From the TkCon, we ran (`>> source write-vel.tcl`). We then added the atomic velocities to the initial `rst7` file with a python script (`>> python add-vels.py`), generating "`v-included.rst7`".

**(Step V2) Generate DMS files on Anton 2.** This step requires access to the Anton 2 supercomputer. We use `viparr` tools to generate Anton-format DMS files separately from (1) our AMBER-format implementation of D.E. Shaw force fields, and (2) official force field files on the Anton 2. For approach 1, we ran the following command on Anton 2:
(`>> viparr-convert-prmtop -c v-included.rst7 -o amber.dms start.prmtop`)
generating DMS file "`amber.dms`". And for approach 2, on Anton 2:
(`>> viparr -f ions.charmm22 -f aa.amber.ff99SB-disp -f water.tip4pd-1.6 -f na.amber.tan2018 amber.dms anton.dms`), generating DMS file "`anton.dms`". The relevant information from both these DMS files was then output to text files, using the following commands on Anton 2: (`>> dms-dump amber.dms > amber.txt`),
(`>> dms-dump anton.dms > anton.txt`).

**(Step V3) Compare Amber-generated and Anton-generated DMS files.** Using the bash script "`run_amber_vs_anton_checks.sh`", we obtained parameters from each of the following: angles, bonds, dihedrals, exclusions, non-bonded, pair, stretch. This script confirmed that the two topologies and all force field parameters matched.

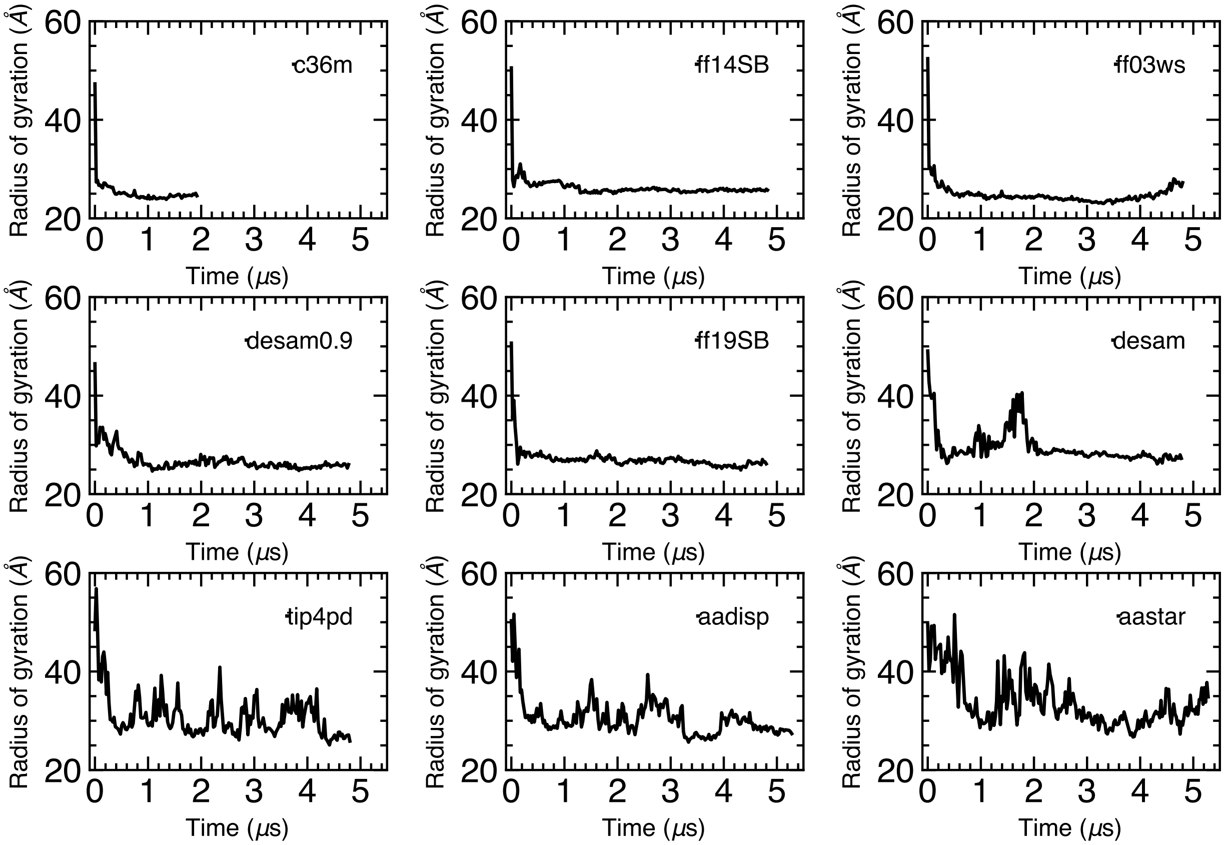


**Figure S1: Radius of gyration of the FUS protein a function of the simulation.** The abbreviated names of the parameter sets are defined in Table 1 in the main text. Data traces for ff14SB, ff03ws, ff19SB, desam, tip4pd, aastar and aadisp are the same as in the main text Fig. 1b and 1c.

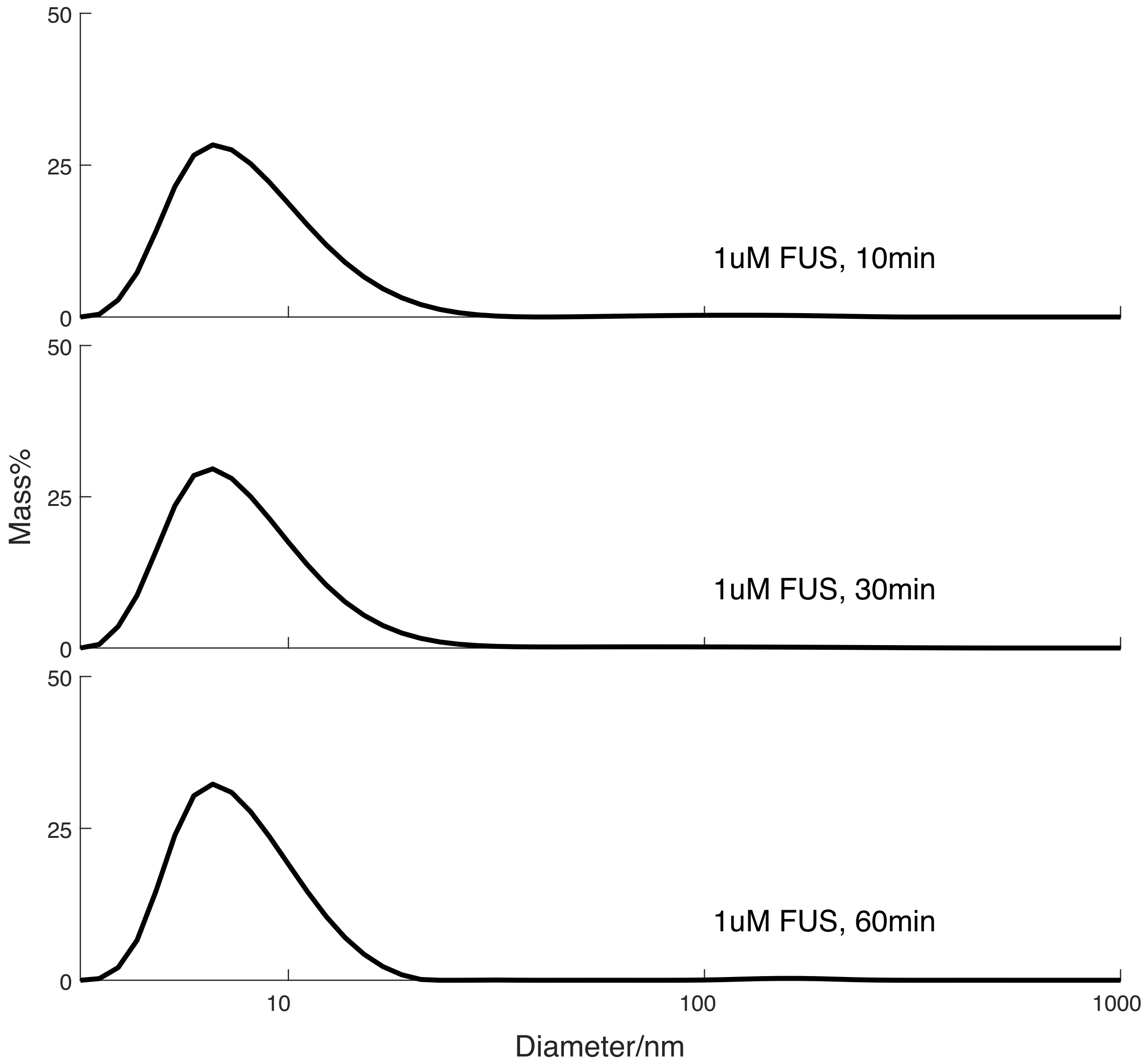


**Figure S2: Time series of DLS measurements of FUS diameter at 1 $\mu$M FUS concentration.** Within 1 hour after the sample was prepared, the diameter distributions centered at around 10 nm. The results in this figure prove the consistency of the size distribution of FUS over time.

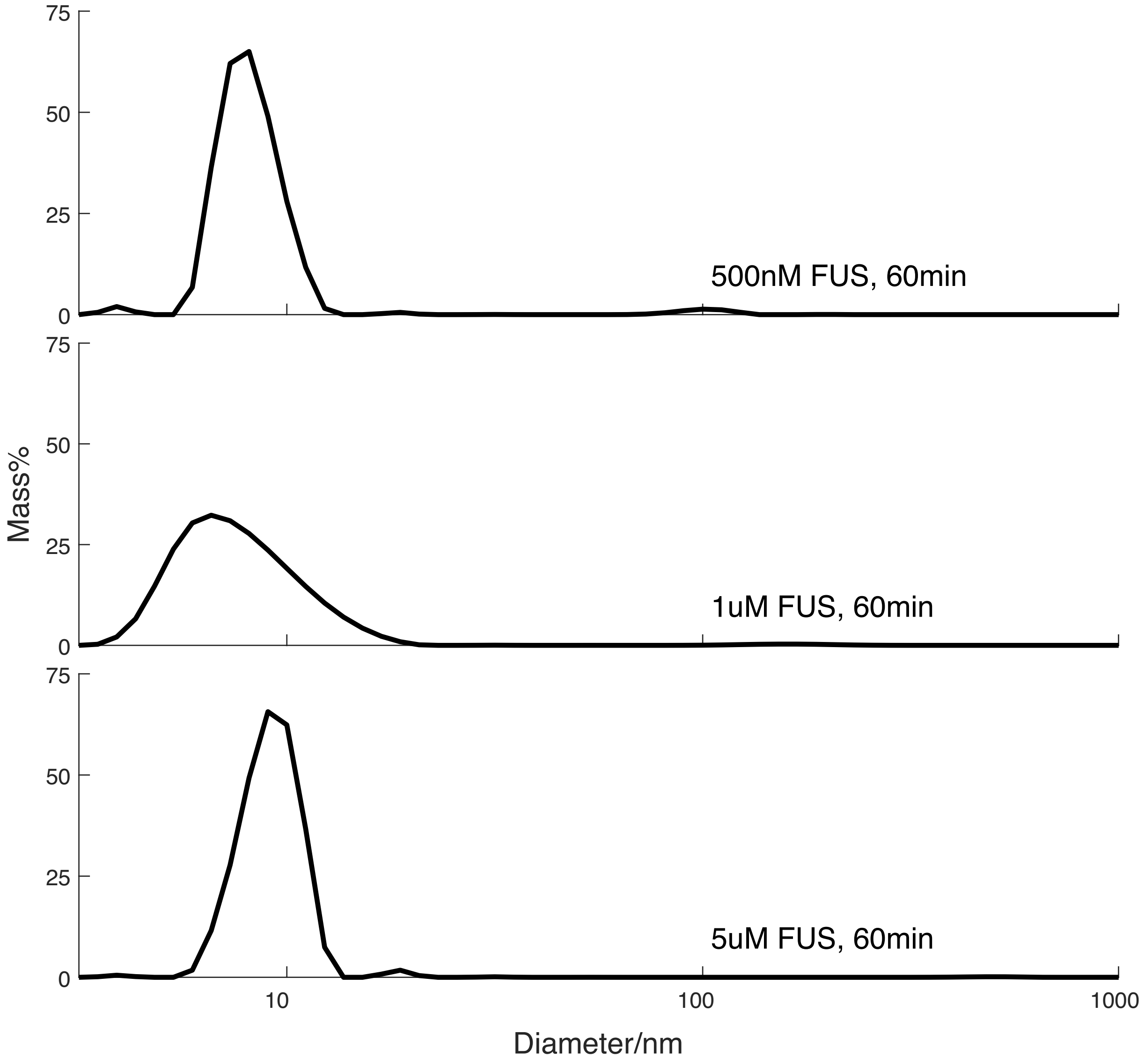


**Figure S3: DLS-measured distributions of FUS diameters at 500 nM, 1 $\mu$M and 5 $\mu$M concentrations of FUS.** The measurements were performed 1 hour after the samples were prepared. For all three of the conditions, the diameter distributions centered at around 10 nm The results in this figure prove that the size distribution of FUS remains constant with the change of concentration.

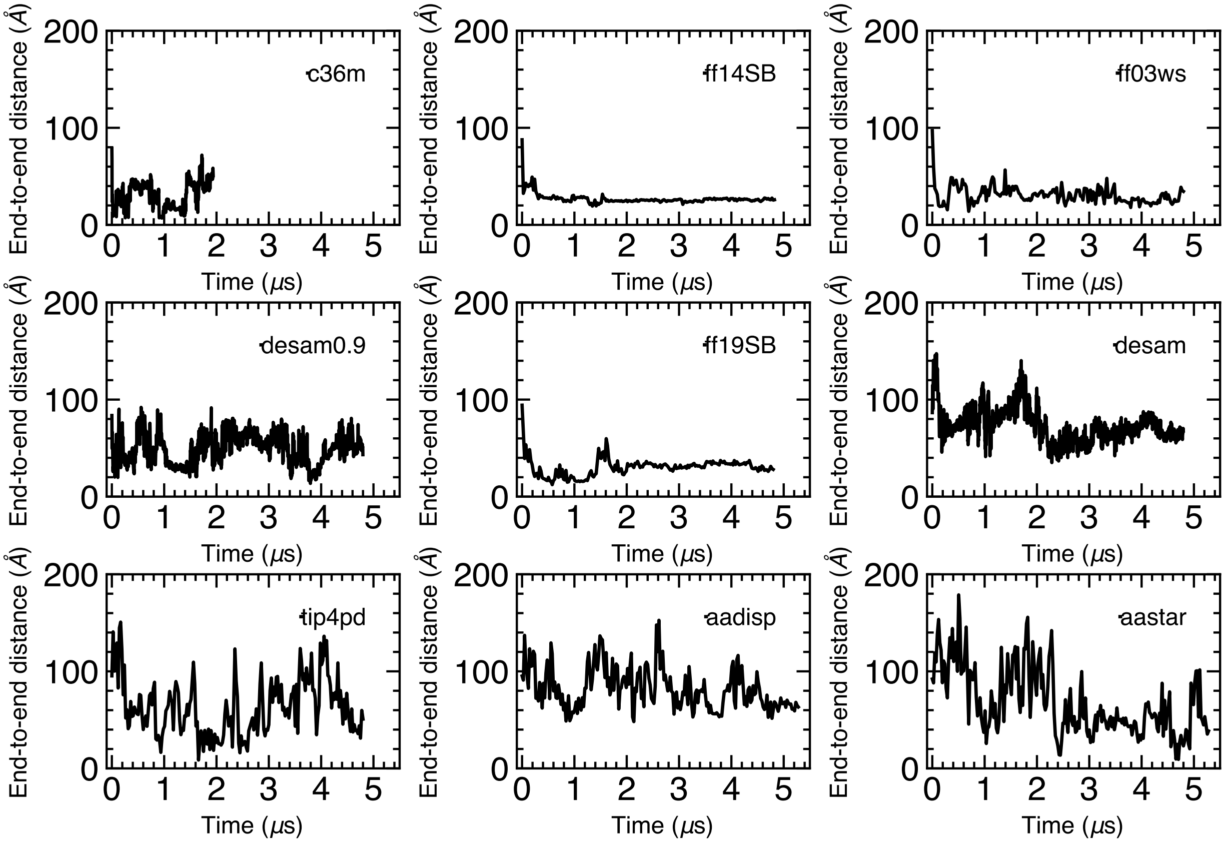


**Figure S4: End to end distance of the FUS protein as a function of the simulation time.** The abbreviated names of the parameter sets are defined in Table 1 in the main text. Data traces for ff14SB, ff03ws, ff19SB, desam, tip4pd, aastar and aadisp are the same as in the main text Fig. 2a and 2b.

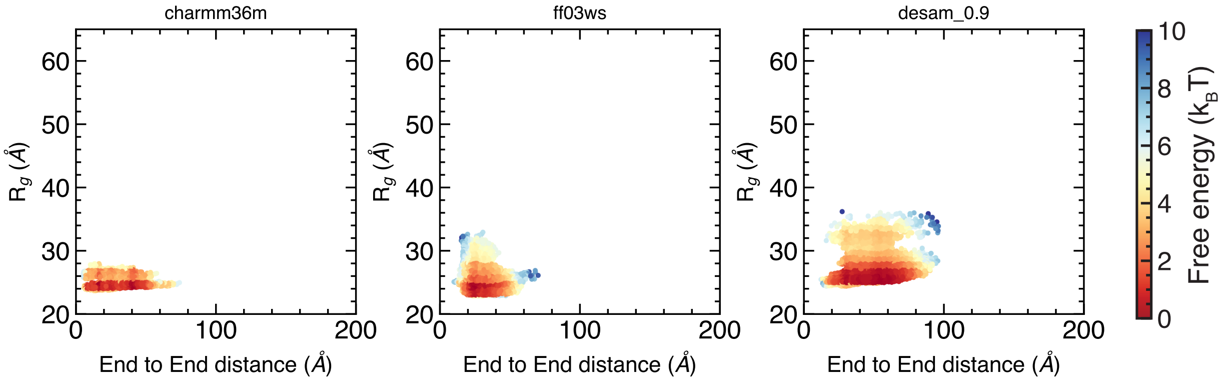


**Figure S5: Free energy map of FUS as a function of its radius of gyration and end-to-end distance.** Each map was constructed by Boltzmann inversion of the conformations sampled by the protein within 5 $\mu$s trajectory, omitting the first 50 ns. The sampling rate was 0.24 ns; the bin size along both coordinates was 1 Å. The reference free energy state for all maps is set at 0, corresponding to a maximum theoretical probability of 1. Given two points on a map, the difference in free energies represents the likelihood of going from one state to the other, favored in the direction of the lower value.

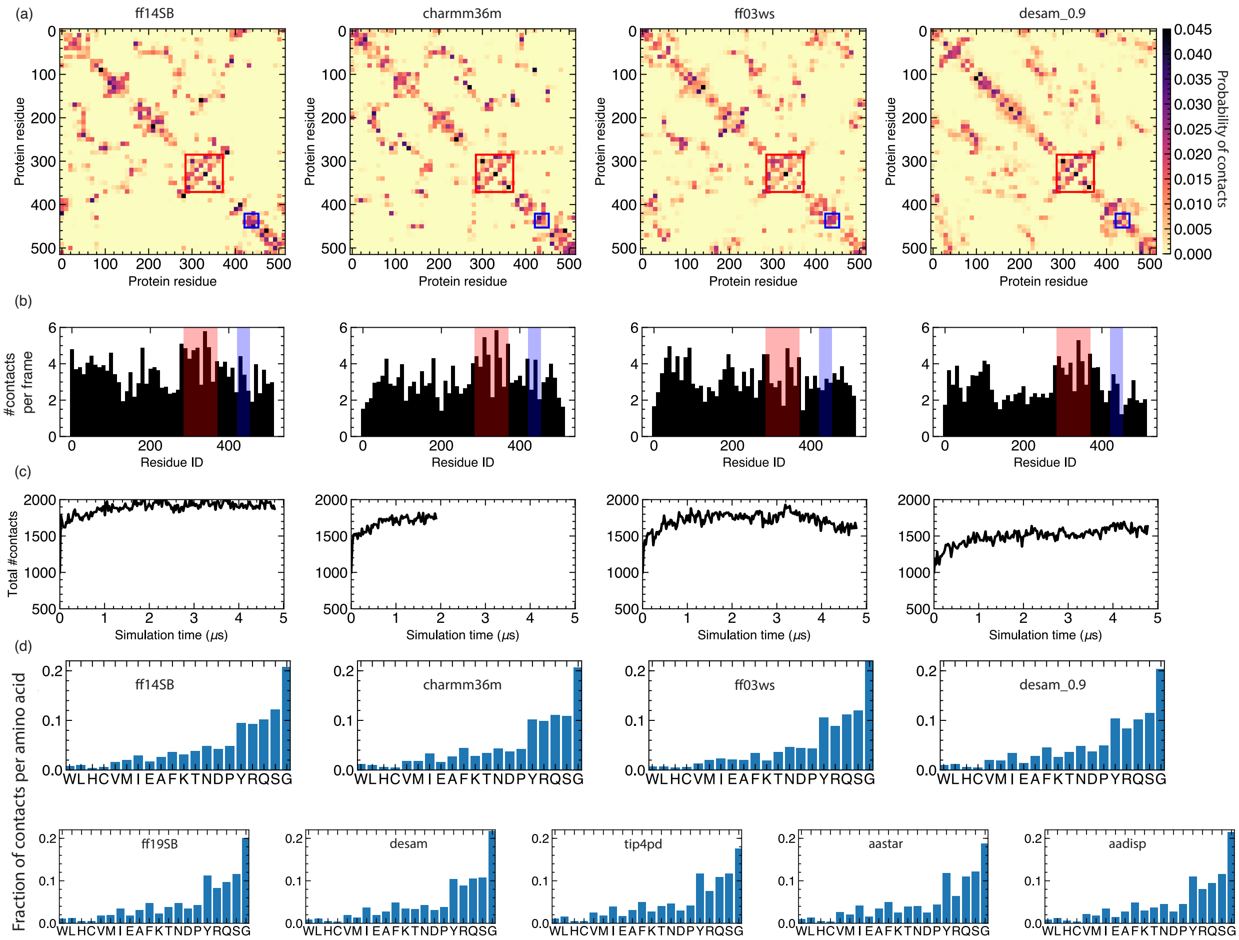


**Figure S6: Intramolecular interactions within FUS.** (a) Residue-based 2D maps of self contacts extracted from the full-length FUS simulation trajectories. Each data point on the 2D map is block averaged over 10 residues in X and Y axis for clarity and the average contact probability is plotted according to the color bar shown on the right. Two residues are considered to be in contact if any atom of one amino acid is located within 3 Å of any atom of the other residue; nearest and next two nearest neighbors are excluded from the analysis. The structured regions of FUS are highlighted in red (RRM) and blue (ZnF). (b) Average number of contacts a given protein residue makes with other residues of FUS (black). Each bar value is block averaged over 10 consecutive residue IDs. The structured regions are highlighted in red (RRM) and blue (ZnF). (c) Total number of unique pairwise intramolecular contacts as a function of simulation time. (d) Trajectory-averaged fraction of contacts formed by a residue of a specified type with other residues of FUS. The amino acids are arranged in ascending order of abundance.

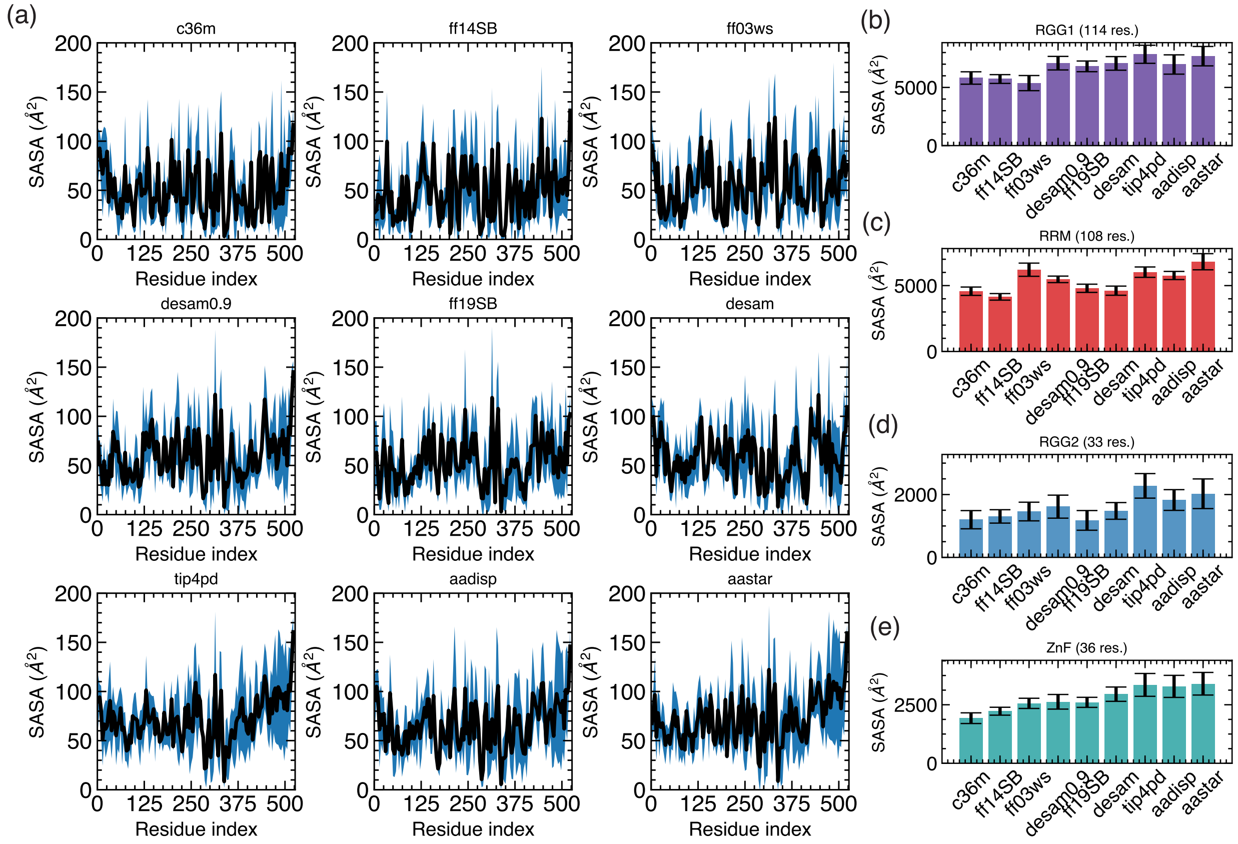


**Figure S7: Solvent accessible surface area.** (a) Solvent accessible surface area (SASA) calculated using VMD[6] for individual FUS residues across the nine parameter sets using a 1.4 Å radius probe, block averaged over 4 FUS residues and time averaged over snapshots taken every 24 ns across 5 $\mu$s of simulation. The error bar represents the standard deviation over the chosen snapshots. The residue SASA values range from 0 to 200 Å, in the right ballpark as expected for connected amino acids in a protein.[7–9] (b-d) SASA calculated for the FUS domains, RGG1 (b), RRM (c), RGG2 (d) and ZnF (e) using a 1.4 Å radius probe, averaged over snapshots taken every 24 ns across 5 $\mu$s of simulation. The error bar represents the standard deviation over the chosen snapshots.

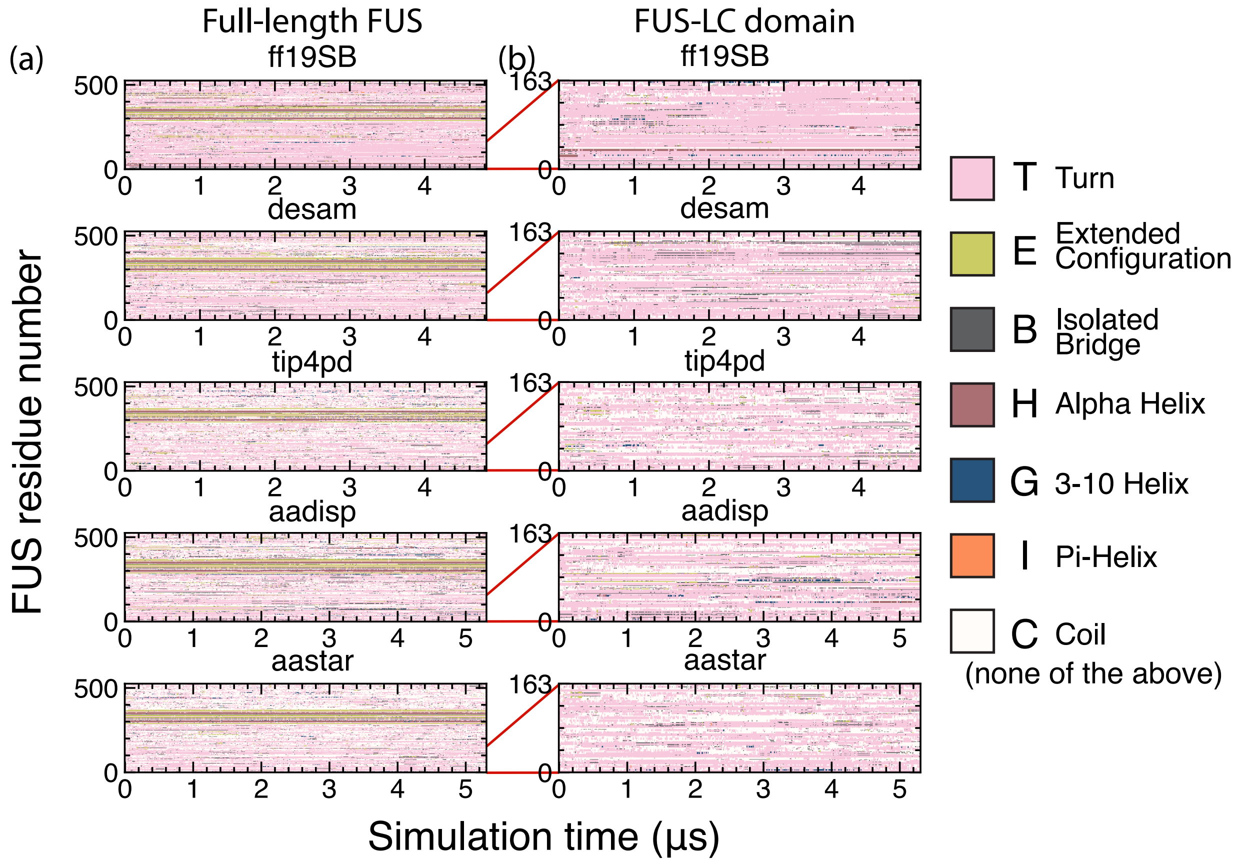


**Figure S8: Secondary structure elements in FUS.** (a) Per-residue secondary structure of the full-length FUS protein (y-axis) as a function of the simulation time for five parameter sets (ff19SB, desam, tip4pd, aadisp and aastar). The secondary structure calculations were done using STRIDE[10] and VMD.[4] (b) A zoomed-in view of the data presented in panel a illustrating secondary structure formation in the LC domain (residues 1 to 163) of FUS. The color key for the secondary structures is shown on the right.

(5) Tan, D.; Piana, S.; Dirks, R. M.; Shaw, D. E. RNA force field with accuracy comparable

to state-of-the-art protein force fields. *Proc. Natl. Acad. Sci. U. S. A.* **2018**, *115*, E1346–E1355.
