## Supplementary material for "Benchmarking Molecular Dynamics Force Fields for All-Atom Simulations of Biological Condensates": Files needed to build FUS in electrolyte: add8.py

```
import parmed
from parmed.tools import addLJType
from parmed.tools import addDihedral
from parmed.amber import AmberParm
import sys

## load prmtop file
PRMTOP=sys.argv[1]
parm = AmberParm(PRMTOP)

## system-specific information
Nresidues = 526
str_n = str(Nresidues)
N_minus_one = Nresidues - 1
str_n_minus_one = str(N_minus_one)

fus1 = "1-526"

fus1_2_to_n_minus_one = ":2-525"
fus1_2_to_n = ":2-526"
fus1_3_to_n = ":3-526"
fus1_1_to_n_minus_one = ":1-525"

f1action1 = addDihedral(parm, fus1_2_to_n_minus_one+"@N", fus1_2_to_n_minus_one+"@CA", fus1_2_to_n_minus_one+"@C", fus1_3_to_n+"@N", -0.197, 1, 180.0)
f1action2 = addDihedral(parm, fus1_2_to_n_minus_one+"@N", fus1_2_to_n_minus_one+"@CA", fus1_2_to_n_minus_one+"@C", fus1_3_to_n+"@N", -0.255, 1, 90.0)
f1action3 = addDihedral(parm, fus1_2_to_n_minus_one+"@N", fus1_2_to_n_minus_one+"@CA", fus1_2_to_n_minus_one+"@C", fus1_3_to_n+"@N",  0.075, 2, 90.0)
f1action4 = addDihedral(parm, fus1_2_to_n_minus_one+"@N", fus1_2_to_n_minus_one+"@CA", fus1_2_to_n_minus_one+"@C", fus1_3_to_n+"@N",  0.057, 3, 90.0)
f1action5 = addDihedral(parm, fus1_2_to_n_minus_one+"@N", fus1_2_to_n_minus_one+"@CA", fus1_2_to_n_minus_one+"@C", fus1_3_to_n+"@N",  0.025, 4, 90.0)
f1action6 = addDihedral(parm, fus1_2_to_n_minus_one+"@N", fus1_2_to_n_minus_one+"@CA", fus1_2_to_n_minus_one+"@C", fus1_3_to_n+"@N", -0.011, 5, 90.0)
f1action8 = addDihedral(parm, fus1_1_to_n_minus_one+"@C", fus1_2_to_n+"@N", fus1_2_to_n+"@CA", fus1_2_to_n+"@C",  0.40, 2, 90.0)

actions = [f1action1, f1action2, f1action3, f1action4, f1action5, f1action6, f1action8]

for i in range(len(actions)):
    actions[i].execute()
    print(str(actions[i]))
parm.save("add8_"+PRMTOP)
```
