## Supplementary material for "Benchmarking Molecular Dynamics Force Fields for All-Atom Simulations of Biological Condensates": Files needed to build FUS in electrolyte: combine_protein_water_ions.sh

```
source leaprc.protein.ffa99SBdisp
source leaprc.water.tip4pew
set default FlexibleWater on

loadoff shaw_fus.lib
loadoff water.lib
loadoff ions.lib
protein_water_ions = combine {shaw_fus,water,ions}
saveamberparm protein_water_ions protein_water_ions.prmtop protein_water_ions.rst7
saveoff protein_water_ions protein_water_ions.lib
quit
```
