## Supplementary material for "Benchmarking Molecular Dynamics Force Fields for All-Atom Simulations of Biological Condensates": Files needed to build FUS in electrolyte: shaw_fus_leaprc.sh

```
source leaprc.RNA.Shaw
source leaprc.protein.ffa99SBdisp.SHAWparm10
source leaprc.water.tip4pew
set default FlexibleWater on

shaw_fus2 = loadpdb "centered_protein.pdb"
saveoff shaw_fus shaw_fus.lib
quit
```
