## Supplementary material for "Benchmarking Molecular Dynamics Force Fields for All-Atom Simulations of Biological Condensates": Files needed to build FUS in electrolyte: water_leaprc.sh

```
source leaprc.protein.ffa99SBdisp
source leaprc.water.tip4pew
set default FlexibleWater on

water = loadpdb "renum_O_only_water.pdb"
saveamberparm water water.prmtop water.rst7
saveoff water water.lib
quit
```
