## Supplementary material for "Benchmarking Molecular Dynamics Force Fields for All-Atom Simulations of Biological Condensates": Files needed to build a small RNA/FUS condensate: add8.py

```
import parmed
from parmed.tools import addLJType
from parmed.tools import addDihedral
from parmed.amber import AmberParm
import sys

## load prmtop file
PRMTOP=sys.argv[1]
parm = AmberParm(PRMTOP)

## system-specific information
Nresidues = 526
str_n = str(Nresidues)
N_minus_one = Nresidues - 1
str_n_minus_one = str(N_minus_one)

fus1 = "1-526"
fus2 = "527-1052"
fus3 = "1053-1578"
fus4 = "1579-2104"
fus5 = "2105-2630"
fus6 = "2631-3156"

fus1_2_to_n_minus_one = ":2-525"
fus1_2_to_n = ":2-526"
fus1_3_to_n = ":3-526"
fus1_1_to_n_minus_one = ":1-525"

fus2_2_to_n_minus_one = ":528-1051"
fus2_2_to_n = ":528-1052"
fus2_3_to_n = ":529-1052"
fus2_1_to_n_minus_one = ":527-1051"

fus3_2_to_n_minus_one = ":1054-1577"
fus3_2_to_n = ":1054-1578"
fus3_3_to_n = ":1055-1578"
fus3_1_to_n_minus_one = ":1053-1577"

fus4_2_to_n_minus_one = ":1580-2103"
fus4_2_to_n = ":1580-2104"
fus4_3_to_n = ":1581-2104"
fus4_1_to_n_minus_one = ":1579-2103"

fus5_2_to_n_minus_one = ":2106-2629"
fus5_2_to_n = ":2106-2630"
fus5_3_to_n = ":2107-2630"
fus5_1_to_n_minus_one = ":2105-2629"

fus6_2_to_n_minus_one = ":2632-3155"
fus6_2_to_n = ":2632-3156"
fus6_3_to_n = ":2633-3156"
fus6_1_to_n_minus_one = ":2631-3155"

# All arguments separate
#action = addLJType(parm, "@1", "radius", 1.5, "epsilon", 0.5)

# Also equivalent; keyword arguments given as keywords
#action = addLJType(parm, "@1", radius=1.5, epsilon=0.5)
## original line:
## action = addDihedral(parm, ":2@N", ":2@CA", ":2@C", ":3@N", -0.255, 1, 90.0)

## Backbone dihedrals to add
## N-CT-C-N (names = N-CA-C-N)
## e.g.
#action1 = addDihedral(parm, ":2-3@N", ":2-3@CA", ":2-3@C", ":3-4@N", -0.197, 1, 180.0)
#action2 = addDihedral(parm, ":2-3@N", ":2-3@CA", ":2-3@C", ":3-4@N", -0.255, 1, 90.0)
#action3 = addDihedral(parm, ":2-3@N", ":2-3@CA", ":2-3@C", ":3-4@N",  0.075, 2, 90.0)
#action4 = addDihedral(parm, ":2-3@N", ":2-3@CA", ":2-3@C", ":3-4@N",  0.057, 3, 90.0)
#action5 = addDihedral(parm, ":2-3@N", ":2-3@CA", ":2-3@C", ":3-4@N",  0.025, 4, 90.0)
#action6 = addDihedral(parm, ":2-3@N", ":2-3@CA", ":2-3@C", ":3-4@N", -0.011, 5, 90.0)

## C-N-CT-C (names = C-N-CA-C)
## e.g.
#action8 = addDihedral(parm, ":1-3@C", ":2-4@N", ":2-4@CA", ":2-4@C",  0.40, 2, 90.0)

f1action1 = addDihedral(parm, fus1_2_to_n_minus_one+"@N", fus1_2_to_n_minus_one+"@CA", fus1_2_to_n_minus_one+"@C", fus1_3_to_n+"@N", -0.197, 1, 180.0)
f1action2 = addDihedral(parm, fus1_2_to_n_minus_one+"@N", fus1_2_to_n_minus_one+"@CA", fus1_2_to_n_minus_one+"@C", fus1_3_to_n+"@N", -0.255, 1, 90.0)
f1action3 = addDihedral(parm, fus1_2_to_n_minus_one+"@N", fus1_2_to_n_minus_one+"@CA", fus1_2_to_n_minus_one+"@C", fus1_3_to_n+"@N",  0.075, 2, 90.0)
f1action4 = addDihedral(parm, fus1_2_to_n_minus_one+"@N", fus1_2_to_n_minus_one+"@CA", fus1_2_to_n_minus_one+"@C", fus1_3_to_n+"@N",  0.057, 3, 90.0)
f1action5 = addDihedral(parm, fus1_2_to_n_minus_one+"@N", fus1_2_to_n_minus_one+"@CA", fus1_2_to_n_minus_one+"@C", fus1_3_to_n+"@N",  0.025, 4, 90.0)
f1action6 = addDihedral(parm, fus1_2_to_n_minus_one+"@N", fus1_2_to_n_minus_one+"@CA", fus1_2_to_n_minus_one+"@C", fus1_3_to_n+"@N", -0.011, 5, 90.0)
f1action8 = addDihedral(parm, fus1_1_to_n_minus_one+"@C", fus1_2_to_n+"@N", fus1_2_to_n+"@CA", fus1_2_to_n+"@C",  0.40, 2, 90.0)

f2action1 = addDihedral(parm, fus2_2_to_n_minus_one+"@N", fus2_2_to_n_minus_one+"@CA", fus2_2_to_n_minus_one+"@C", fus2_3_to_n+"@N", -0.197, 1, 180.0)
f2action2 = addDihedral(parm, fus2_2_to_n_minus_one+"@N", fus2_2_to_n_minus_one+"@CA", fus2_2_to_n_minus_one+"@C", fus2_3_to_n+"@N", -0.255, 1, 90.0)
f2action3 = addDihedral(parm, fus2_2_to_n_minus_one+"@N", fus2_2_to_n_minus_one+"@CA", fus2_2_to_n_minus_one+"@C", fus2_3_to_n+"@N",  0.075, 2, 90.0)
f2action4 = addDihedral(parm, fus2_2_to_n_minus_one+"@N", fus2_2_to_n_minus_one+"@CA", fus2_2_to_n_minus_one+"@C", fus2_3_to_n+"@N",  0.057, 3, 90.0)
f2action5 = addDihedral(parm, fus2_2_to_n_minus_one+"@N", fus2_2_to_n_minus_one+"@CA", fus2_2_to_n_minus_one+"@C", fus2_3_to_n+"@N",  0.025, 4, 90.0)
f2action6 = addDihedral(parm, fus2_2_to_n_minus_one+"@N", fus2_2_to_n_minus_one+"@CA", fus2_2_to_n_minus_one+"@C", fus2_3_to_n+"@N", -0.011, 5, 90.0)
f2action8 = addDihedral(parm, fus2_1_to_n_minus_one+"@C", fus2_2_to_n+"@N", fus2_2_to_n+"@CA", fus2_2_to_n+"@C",  0.40, 2, 90.0)

f3action1 = addDihedral(parm, fus3_2_to_n_minus_one+"@N", fus3_2_to_n_minus_one+"@CA", fus3_2_to_n_minus_one+"@C", fus3_3_to_n+"@N", -0.197, 1, 180.0)
f3action2 = addDihedral(parm, fus3_2_to_n_minus_one+"@N", fus3_2_to_n_minus_one+"@CA", fus3_2_to_n_minus_one+"@C", fus3_3_to_n+"@N", -0.255, 1, 90.0)
f3action3 = addDihedral(parm, fus3_2_to_n_minus_one+"@N", fus3_2_to_n_minus_one+"@CA", fus3_2_to_n_minus_one+"@C", fus3_3_to_n+"@N",  0.075, 2, 90.0)
f3action4 = addDihedral(parm, fus3_2_to_n_minus_one+"@N", fus3_2_to_n_minus_one+"@CA", fus3_2_to_n_minus_one+"@C", fus3_3_to_n+"@N",  0.057, 3, 90.0)
f3action5 = addDihedral(parm, fus3_2_to_n_minus_one+"@N", fus3_2_to_n_minus_one+"@CA", fus3_2_to_n_minus_one+"@C", fus3_3_to_n+"@N",  0.025, 4, 90.0)
f3action6 = addDihedral(parm, fus3_2_to_n_minus_one+"@N", fus3_2_to_n_minus_one+"@CA", fus3_2_to_n_minus_one+"@C", fus3_3_to_n+"@N", -0.011, 5, 90.0)
f3action8 = addDihedral(parm, fus3_1_to_n_minus_one+"@C", fus3_2_to_n+"@N", fus3_2_to_n+"@CA", fus3_2_to_n+"@C",  0.40, 2, 90.0)

f4action1 = addDihedral(parm, fus4_2_to_n_minus_one+"@N", fus4_2_to_n_minus_one+"@CA", fus4_2_to_n_minus_one+"@C", fus4_3_to_n+"@N", -0.197, 1, 180.0)
f4action2 = addDihedral(parm, fus4_2_to_n_minus_one+"@N", fus4_2_to_n_minus_one+"@CA", fus4_2_to_n_minus_one+"@C", fus4_3_to_n+"@N", -0.255, 1, 90.0)
f4action3 = addDihedral(parm, fus4_2_to_n_minus_one+"@N", fus4_2_to_n_minus_one+"@CA", fus4_2_to_n_minus_one+"@C", fus4_3_to_n+"@N",  0.075, 2, 90.0)
f4action4 = addDihedral(parm, fus4_2_to_n_minus_one+"@N", fus4_2_to_n_minus_one+"@CA", fus4_2_to_n_minus_one+"@C", fus4_3_to_n+"@N",  0.057, 3, 90.0)
f4action5 = addDihedral(parm, fus4_2_to_n_minus_one+"@N", fus4_2_to_n_minus_one+"@CA", fus4_2_to_n_minus_one+"@C", fus4_3_to_n+"@N",  0.025, 4, 90.0)
f4action6 = addDihedral(parm, fus4_2_to_n_minus_one+"@N", fus4_2_to_n_minus_one+"@CA", fus4_2_to_n_minus_one+"@C", fus4_3_to_n+"@N", -0.011, 5, 90.0)
f4action8 = addDihedral(parm, fus4_1_to_n_minus_one+"@C", fus4_2_to_n+"@N", fus4_2_to_n+"@CA", fus4_2_to_n+"@C",  0.40, 2, 90.0)

f5action1 = addDihedral(parm, fus5_2_to_n_minus_one+"@N", fus5_2_to_n_minus_one+"@CA", fus5_2_to_n_minus_one+"@C", fus5_3_to_n+"@N", -0.197, 1, 180.0)
f5action2 = addDihedral(parm, fus5_2_to_n_minus_one+"@N", fus5_2_to_n_minus_one+"@CA", fus5_2_to_n_minus_one+"@C", fus5_3_to_n+"@N", -0.255, 1, 90.0)
f5action3 = addDihedral(parm, fus5_2_to_n_minus_one+"@N", fus5_2_to_n_minus_one+"@CA", fus5_2_to_n_minus_one+"@C", fus5_3_to_n+"@N",  0.075, 2, 90.0)
f5action4 = addDihedral(parm, fus5_2_to_n_minus_one+"@N", fus5_2_to_n_minus_one+"@CA", fus5_2_to_n_minus_one+"@C", fus5_3_to_n+"@N",  0.057, 3, 90.0)
f5action5 = addDihedral(parm, fus5_2_to_n_minus_one+"@N", fus5_2_to_n_minus_one+"@CA", fus5_2_to_n_minus_one+"@C", fus5_3_to_n+"@N",  0.025, 4, 90.0)
f5action6 = addDihedral(parm, fus5_2_to_n_minus_one+"@N", fus5_2_to_n_minus_one+"@CA", fus5_2_to_n_minus_one+"@C", fus5_3_to_n+"@N", -0.011, 5, 90.0)
f5action8 = addDihedral(parm, fus5_1_to_n_minus_one+"@C", fus5_2_to_n+"@N", fus5_2_to_n+"@CA", fus5_2_to_n+"@C",  0.40, 2, 90.0)

f6action1 = addDihedral(parm, fus6_2_to_n_minus_one+"@N", fus6_2_to_n_minus_one+"@CA", fus6_2_to_n_minus_one+"@C", fus6_3_to_n+"@N", -0.197, 1, 180.0)
f6action2 = addDihedral(parm, fus6_2_to_n_minus_one+"@N", fus6_2_to_n_minus_one+"@CA", fus6_2_to_n_minus_one+"@C", fus6_3_to_n+"@N", -0.255, 1, 90.0)
f6action3 = addDihedral(parm, fus6_2_to_n_minus_one+"@N", fus6_2_to_n_minus_one+"@CA", fus6_2_to_n_minus_one+"@C", fus6_3_to_n+"@N",  0.075, 2, 90.0)
f6action4 = addDihedral(parm, fus6_2_to_n_minus_one+"@N", fus6_2_to_n_minus_one+"@CA", fus6_2_to_n_minus_one+"@C", fus6_3_to_n+"@N",  0.057, 3, 90.0)
f6action5 = addDihedral(parm, fus6_2_to_n_minus_one+"@N", fus6_2_to_n_minus_one+"@CA", fus6_2_to_n_minus_one+"@C", fus6_3_to_n+"@N",  0.025, 4, 90.0)
f6action6 = addDihedral(parm, fus6_2_to_n_minus_one+"@N", fus6_2_to_n_minus_one+"@CA", fus6_2_to_n_minus_one+"@C", fus6_3_to_n+"@N", -0.011, 5, 90.0)
f6action8 = addDihedral(parm, fus6_1_to_n_minus_one+"@C", fus6_2_to_n+"@N", fus6_2_to_n+"@CA", fus6_2_to_n+"@C",  0.40, 2, 90.0)

actions = [f1action1, f1action2, f1action3, f1action4, f1action5, f1action6, f1action8,
           f2action1, f2action2, f2action3, f2action4, f2action5, f2action6, f2action8,
           f3action1, f3action2, f3action3, f3action4, f3action5, f3action6, f3action8,
           f4action1, f4action2, f4action3, f4action4, f4action5, f4action6, f4action8,
           f5action1, f5action2, f5action3, f5action4, f5action5, f5action6, f5action8,
           f6action1, f6action2, f6action3, f6action4, f6action5, f6action6, f6action8]

for i in range(len(actions)):
    actions[i].execute()
    print(str(actions[i]))
parm.save("add8_"+PRMTOP)
```
