## Supplementary material for "Benchmarking Molecular Dynamics Force Fields for All-Atom Simulations of Biological Condensates": Files needed to build a small RNA/FUS condensate: combine_protein_rna_water_ions.py

```
import time
import parmed as pmd

## for timing
prefix = "protein_rna_water_ions"
start_time = time.time()

parm_protein_rna = pmd.load_file('protein_rna.prmtop', 'protein_rna.rst7')
parm_water = pmd.load_file('reoriented_water.prmtop', 'reoriented_water.rst7')
parm_ions = pmd.load_file('ions.prmtop', 'ions.rst7')

## output time to load
dt_seconds_loading = ( time.time() - start_time )
dt_minutes_loading = dt_seconds_loading/(60.0)
outfile_loading = open(prefix + ".loading.dat", "w")
outfile_loading.write("# Natoms, minutes" + "\n")
outfile_loading.write(str(dt_minutes_loading) + "\n")
outfile_loading.close()

## combining
protein_rna_water_ions = parm_protein_rna + parm_water + parm_ions
protein_rna_water_ions.save('protein_rna_water_ions.prmtop')
protein_rna_water_ions.save('protein_rna_water_ions.rst7')

## output the time it took
dt_seconds = ( time.time() - start_time )
dt_minutes = dt_seconds/(60.0)
outfile_timing = open(prefix + ".timing.dat", "w")
outfile_timing.write("# Natoms, minutes" + "\n")
outfile_timing.write(str(dt_minutes) + "\n")
outfile_timing.close()
```
