## Supplementary material for "Benchmarking Molecular Dynamics Force Fields for All-Atom Simulations of Biological Condensates": Files needed to build a small RNA/FUS condensate: shaw_combine_protein_rna_water_ions.sh

```
#source /data/server1/dwinogra/fromHimanshu/amber16/dat/leap/cmd/leaprc.RNA.OL3
source leaprc.RNA.Shaw
source leaprc.protein.ffa99SBdisp.SHAWparm10
source leaprc.water.tip4pew
set default FlexibleWater on

loadoff shaw_protein.lib
loadoff shaw_rna_x3.lib
loadoff shaw_water.lib
loadoff shaw_ions.lib
shaw_protein_rna_water_ions = combine {shaw_protein,shaw_rna_x3,shaw_water,shaw_ions}
saveamberparm shaw_protein_rna_water_ions shaw_protein_rna_water_ions.prmtop shaw_protein_rna_water_ions.rst7
saveoff shaw_protein_rna_water_ions.lib
quit
```
