## Supplementary material for "Benchmarking Molecular Dynamics Force Fields for All-Atom Simulations of Biological Condensates": Files needed to build a small RNA/FUS condensate: ions_readpdb.py

```
import numpy as np 
import sys
f = open('renum2_ions_fus64_310K_amberNamed.pdb','w')
f2 = open('renum_ions_fus64_310K_amberNamed.pdb','r')
lines2 = f2.readlines()
lines2 = lines2[1:-2]
f2.close()

widths = (0,6,5,5,5,2,7,8,8,8,6,6,8,6)
field_start = np.cumsum(widths)[:-1]
field_end = np.cumsum(widths)[1:]
i=0
for i in range(len(lines2)):
    print(i) ## for debugging
    l2 = [lines2[i][a:b] for a,b in zip(field_start,field_end)]
    #print(len(l2), l2)
    #length_l2=len(l2)
    i=i+1
    f.write("{:6s}{:5s}{:^4s}{:3s}{:<8d} {:8.3f}{:8.3f}{:8.3f}{:6.2f}{:6.2f}       {:1s}  {:4s}".format("ATOM",l2[1],l2[2],l2[3],i,float(l2[6]),float(l2[7]),float(l2[8]),float(l2[9]),float(l2[10]),'A',l2[11]))
    f.write("\n")
f.close()
```
