## Supplementary material for "Benchmarking Molecular Dynamics Force Fields for All-Atom Simulations of Biological Condensates": Files needed to build a small RNA/FUS condensate: shaw_ions_leaprc.sh

```
#source /data/server1/dwinogra/fromHimanshu/amber16/dat/leap/cmd/leaprc.RNA.OL3
#source leaprc.protein.ffa99SBdisp.parm10
source leaprc.RNA.Shaw
source leaprc.protein.ffa99SBdisp.SHAWparm10
source leaprc.water.tip4pew
set default FlexibleWater on

shaw_ions = loadpdb "ions.with_RNA.pdb"
saveamberparm shaw_ions shaw_ions.prmtop shaw_ions.rst7
saveoff shaw_ions shaw_ions.lib
quit
```
