## Supplementary material for "Benchmarking Molecular Dynamics Force Fields for All-Atom Simulations of Biological Condensates": Files needed to build a small RNA/FUS condensate: combine_3rna_leaprc.sh

```
source leaprc.RNA.Shaw
source leaprc.water.tip4pew
set default FlexibleWater on

loadoff shaw_rna1.lib
loadoff shaw_rna2.lib
loadoff shaw_rna3.lib
shaw_rna_x3 = combine {shaw_rna1,shaw_rna2,shaw_rna3}
saveoff shaw_rna_x3 shaw_rna_x3.lib
saveamberparm shaw_rna_x3 shaw_rna_x3.prmtop shaw_rna_x3.rst7
quit
```
