## Supplementary material for "Benchmarking Molecular Dynamics Force Fields for All-Atom Simulations of Biological Condensates": Files needed to build a small RNA/FUS condensate: june29_write_separate_nucleic_files.tcl

```
set inPrefix noh_nucleic.with_RNA
mol new $inPrefix.pdb
set all [atomselect top "all"]
set segNames [lsort -unique [$all get segname]]
set COUNTER 1
foreach seg $segNames {
    set segi [atomselect top "segname $seg"]
    $segi writepdb june29_rna$COUNTER.pdb
    $segi delete
    set $COUNTER [expr {$COUNTER+1}]
}
exit
```
