## Supplementary material for "Benchmarking Molecular Dynamics Force Fields for All-Atom Simulations of Biological Condensates": Files needed to build a small RNA/FUS condensate: rna_x3_leaprc.sh

```
source /data/server1/dwinogra/fromHimanshu/amber16/dat/leap/cmd/leaprc.RNA.OL3
#source leaprc.protein.ffa99SBdisp.parm10
source leaprc.water.tip4pew
set default FlexibleWater on

rna1 = loadpdb "renum_rna1_R.pdb"
saveamberparm rna1 rna1.prmtop rna1.rst7
rna2 = loadpdb "renum_rna2_R.pdb"
saveamberparm rna2 rna2.prmtop rna2.rst7
rna3 = loadpdb "renum_rna3_R.pdb"
saveamberparm rna3 rna3.prmtop rna3.rst7
quit
```
