## Supplementary material for "Benchmarking Molecular Dynamics Force Fields for All-Atom Simulations of Biological Condensates": Files needed to build a small RNA/FUS condensate: shaw_rna_x3_leaprc.sh

```
source leaprc.RNA.Shaw
source leaprc.water.tip4pew
set default FlexibleWater on

shaw_rna1 = loadpdb "renum_rna1_R.pdb"
saveoff shaw_rna1 shaw_rna1.lib
saveamberparm shaw_rna1 shaw_rna1.prmtop shaw_rna1.rst7
shaw_rna2 = loadpdb "renum_rna2_R.pdb"
saveoff shaw_rna2 shaw_rna2.lib
saveamberparm shaw_rna2 shaw_rna2.prmtop shaw_rna2.rst7
shaw_rna3 = loadpdb "renum_rna3_R.pdb"
saveoff shaw_rna3 shaw_rna3.lib
saveamberparm shaw_rna3 shaw_rna3.prmtop shaw_rna3.rst7

quit
```
