## Supplementary material for "Benchmarking Molecular Dynamics Force Fields for All-Atom Simulations of Biological Condensates": Files needed to build a small RNA/FUS condensate: combine_6fus_leaprc.sh

```
source /data/server1/dwinogra/fromHimanshu/amber16/dat/leap/cmd/leaprc.RNA.OL3
source leaprc.protein.ffa99SBdisp.parm10
source leaprc.water.tip4pew
set default FlexibleWater on

loadoff fus1.lib
loadoff fus2.lib
loadoff fus3.lib
loadoff fus4.lib
loadoff fus5.lib
loadoff fus6.lib
protein = combine {fus1,fus2,fus3,fus4,fus5,fus6}
saveamberparm protein protein.prmtop protein.rst7
quit
```
