## Supplementary material for "Benchmarking Molecular Dynamics Force Fields for All-Atom Simulations of Biological Condensates": Files needed to build a small RNA/FUS condensate: fus_x6_leaprc.sh

```
source /data/server1/dwinogra/fromHimanshu/amber16/dat/leap/cmd/leaprc.RNA.OL3
source leaprc.protein.ffa99SBdisp.parm10
source leaprc.water.tip4pew
set default FlexibleWater on

fus1 = loadpdb "fus.1.pdb"
saveoff fus1 fus1.lib
fus2 = loadpdb "fus.2.pdb"
saveoff fus2 fus2.lib
fus3 = loadpdb "fus.3.pdb"
saveoff fus3 fus3.lib
fus4 = loadpdb "fus.4.pdb"
saveoff fus4 fus4.lib
fus5 = loadpdb "fus.5.pdb"
saveoff fus5 fus5.lib
fus6 = loadpdb "fus.6.pdb"
saveoff fus6 fus6.lib
quit
```
