## Supplementary material for "Benchmarking Molecular Dynamics Force Fields for All-Atom Simulations of Biological Condensates": Files needed to build a small RNA/FUS condensate: shaw_combine_6fus_leaprc.sh

```
#source /data/server1/dwinogra/fromHimanshu/amber16/dat/leap/cmd/leaprc.RNA.OL3
source leaprc.RNA.Shaw
source leaprc.protein.ffa99SBdisp.SHAWparm10
source leaprc.water.tip4pew
set default FlexibleWater on

loadoff shaw_fus1.lib
loadoff shaw_fus2.lib
loadoff shaw_fus3.lib
loadoff shaw_fus4.lib
loadoff shaw_fus5.lib
loadoff shaw_fus6.lib
shaw_protein = combine {shaw_fus1,shaw_fus2,shaw_fus3,shaw_fus4,shaw_fus5,shaw_fus6}
saveamberparm shaw_protein shaw_protein.prmtop shaw_protein.rst7
saveoff shaw_protein shaw_protein.lib
quit
```
