## Supplementary material for "Benchmarking Molecular Dynamics Force Fields for All-Atom Simulations of Biological Condensates": Files needed to build a small RNA/FUS condensate: shaw_fus_x6_leaprc.sh

```
#source /data/server1/dwinogra/fromHimanshu/amber16/dat/leap/cmd/leaprc.RNA.OL3
source leaprc.RNA.Shaw
source leaprc.protein.ffa99SBdisp.SHAWparm10
source leaprc.water.tip4pew
set default FlexibleWater on

shaw_fus1 = loadpdb "fus.1.pdb"
saveoff shaw_fus1 shaw_fus1.lib
shaw_fus2 = loadpdb "fus.2.pdb"
saveoff shaw_fus2 shaw_fus2.lib
shaw_fus3 = loadpdb "fus.3.pdb"
saveoff shaw_fus3 shaw_fus3.lib
shaw_fus4 = loadpdb "fus.4.pdb"
saveoff shaw_fus4 shaw_fus4.lib
shaw_fus5 = loadpdb "fus.5.pdb"
saveoff shaw_fus5 shaw_fus5.lib
shaw_fus6 = loadpdb "fus.6.pdb"
saveoff shaw_fus6 shaw_fus6.lib
quit
```
