## Supplementary material for "Benchmarking Molecular Dynamics Force Fields for All-Atom Simulations of Biological Condensates": Files needed to build a small RNA/FUS condensate: looped_readpdb.py

```
import numpy as np 
import sys

prefix = "noh_water_fus64_310K_amberNamed"
for ii in range(52):
    print("FILE ", ii)
    filename_ii = prefix + "." + str(ii) + ".pdb"
    f = open('renum_'+filename_ii,'w')
    f2 = open(filename_ii,'r')
    lines2 = f2.readlines()
    lines2 = lines2[1:-2]
    f2.close()
    
    widths = (0,6,5,5,5,2,7,8,8,8,6,6,8,6)
    field_start = np.cumsum(widths)[:-1]
    field_end = np.cumsum(widths)[1:]
    
    i=0
    for i in range(len(lines2)):
        print(i) ## for debugging
        l2 = [lines2[i][a:b] for a,b in zip(field_start,field_end)]
        #print(len(l2), l2)
        #length_l2=len(l2)
        i=i+1
        f.write("{:6s}{:5s}{:^4s}{:3s}{:<8d} {:8.3f}{:8.3f}{:8.3f}{:6.2f}{:6.2f}       {:1s}  {:4s}".format("ATOM",l2[1],l2[2],l2[3],i,float(l2[6]),float(l2[7]),float(l2[8]),float(l2[9]),float(l2[10]),'A',l2[11]))
        f.write("\n")
    f.close()
```
