## Supplementary material for "Benchmarking Molecular Dynamics Force Fields for All-Atom Simulations of Biological Condensates": Files needed to build a small RNA/FUS condensate: orient_waters.tcl

```
set prefix water
mol new $prefix.prmtop
mol addfile $prefix.rst7
set all0 [atomselect 0 "all"]
set three_points_0 [atomselect 0 "not name EPW"]
set residues0 [lsort -unique [$all0 get residue]]

mol new Hincluded_water.residue10plus.pdb
set all1 [atomselect 1 "all"]
set residues1 [lsort -unique [$all1 get residue]]

set COUNTER 1
foreach r0 $residues0 r1 $residues1 {
    set sel1 [atomselect 1 "residue $r1"]
    set O_1 [atomselect 1 "residue $r1 and (name O)"]
    set com1 [measure center $O_1]
    set minus1 [vecscale -1 $com1]
    $sel1 moveby $minus1
    
    set sel0 [atomselect 0 "residue $r0"]
    set O_0 [atomselect 0 "residue $r0 and (name O)"]
    set three_point_0 [atomselect 0 "not name EPW and (residue $r0)"]
    set com0 [measure center $O_0]
    set minus0 [vecscale -1 $com0]
    $sel0 moveby $minus0
    
    set M [measure fit $three_point_0 $sel1]
    $sel0 move $M
    $sel0 moveby $com1
    $sel1 moveby $com1
    
    $sel1 delete
    $sel0 delete
    $three_point_0 delete
    $O_0 delete
    $O_1 delete
    
    puts "$COUNTER waters moved"
    set COUNTER [expr {$COUNTER + 1}]
}

puts "waters reoriented :)"
```
