## Supplementary material for "Benchmarking Molecular Dynamics Force Fields for All-Atom Simulations of Biological Condensates": Files needed to build a small RNA/FUS condensate: shaw_water_redone_leaprc.sh

```
source leaprc.RNA.Shaw
source leaprc.protein.ffa99SBdisp.SHAWparm10
source leaprc.water.tip4pew
set default FlexibleWater on

shaw_water = loadpdb "renum_noh_water.with_RNA.pdb"
saveamberparm shaw_water shaw_water_redone.prmtop shaw_water_redone.rst7
saveoff shaw_water shaw_water.lib
quit
```
