## Supplementary material for "Benchmarking Molecular Dynamics Force Fields for All-Atom Simulations of Biological Condensates": Files needed to build a small RNA/FUS condensate: water_redone_leaprc.sh

```
source /data/server1/dwinogra/fromHimanshu/amber16/dat/leap/cmd/leaprc.RNA.OL3
source leaprc.protein.ffa99SBdisp.parm10
source leaprc.water.tip4pew
set default FlexibleWater on

water = loadpdb "renum_noh_water.with_RNA.pdb"
saveamberparm water water_redone.prmtop water_redone.rst7
#saveoff water water.lib
quit
```
