## Supplementary material for "Benchmarking Molecular Dynamics Force Fields for All-Atom Simulations of Biological Condensates": Files needed to build a small RNA/FUS condensate: add-vels.py

```
import numpy as np

a = open('namd_vels_124940.txt','r')
lines = a.readlines()
a.close()

a = open('centered_add8_shaw_protein_rna_water_ions.rst7','r') #original rst7
l1 = a.readlines()
a.close()

f = open('vincluded_centered_add8_shaw_protein_rna_water_ions.rst7','w') #New rst7

total_string = ""

print("adding velocities...")
for i in range(len(lines)):
    a1 = lines[i].split()
    a2 = float(a1[0])
    a3 = float(a1[1])
    a4 = float(a1[2])
    string = " {:11.7f} {:11.7f} {:11.7f}".format(a2,a3,a4)
    k = ""
    if i%2==1:
        k = "\n"
    string = string + k
    total_string = total_string + string
total_string = total_string + '\n'

for i in range(len(l1)-1):
    f.write(l1[i])
f.write(total_string)
f.write(l1[len(l1)-1])
f.close()
print("Done :)")
```
