## Supplementary material for "Benchmarking Molecular Dynamics Force Fields for All-Atom Simulations of Biological Condensates": Archive of leap files: Makefile

```
all:  ff91 ff94 ff02 ff02EP ff03ua ff03 ff10 ions91 ions94 ions08 solvents

ff91: 
	/bin/rm -f ff91.log
	../../../bin/teLeap -s -f ff91.cmd

ff94: 
	/bin/rm -f ff94.log
	../../../bin/teLeap -s -f ff94.cmd

ff02: 
	/bin/rm -f ff02.log
	../../../bin/teLeap -s -f ff02.cmd

ff02EP: 
	/bin/rm -f ff02EP.log
	../../../bin/teLeap -s -f ff02EP.cmd

ff03ua: 
	/bin/rm -f ff03ua.log
	../../../bin/teLeap -s -f ff03ua.cmd

ions91: 
	/bin/rm -f ions91.log
	../../../bin/teLeap -s -f ions91.cmd

ions94: 
	/bin/rm -f ions91.log
	../../../bin/teLeap -s -f ions94.cmd

ions08: 
	/bin/rm -f ions08.log
	../../../bin/teLeap -s -f ions08.cmd

solvents:
	/bin/rm -f solvents.log
	../../../bin/teLeap -s -f solvents.cmd

ff03:
	/bin/rm -f ff03.log
	../../../bin/teLeap -s -f ff03.cmd

ff10:
	/bin/rm -f ff10.log
	../../../bin/teLeap -s -f ff10.cmd

ff12polL:
	/bin/rm -f ff12polL.log
	../../../bin/teLeap -s -f ff12polL.cmd

ff12polA:
	/bin/rm -f ff12polA.log
	../../../bin/teLeap -s -f ff12polA.cmd
```
