## Supplementary material for "Benchmarking Molecular Dynamics Force Fields for All-Atom Simulations of Biological Condensates": Archive of leap files: validate_torsions.py

```
"""
This script will validate the torsions specified in a particular database and
will highlight any specific torsions that need additional terms to explicitly
override generic torsions
"""
import os
import parmed as pmd
import sys

if len(sys.argv) < 2:
    sys.exit('%s <param_file> [<param_file> [<param_file> ...]]' %
             os.path.split(sys.argv[0])[1])

params = pmd.amber.AmberParameterSet(sys.argv[1:])

# First separate the generic terms and the specific terms. Store the generics as
# *just* the middle terms, and do both permutations
generics = dict()
generic_type_ids = set()
specifics = dict()
specific_type_ids = set()

for key, dihtype in params.dihedral_types.items():
    if key[0] == 'X' and key[3] == 'X':
        if id(dihtype) in generic_type_ids:
            continue
        generics[(key[1], key[2])] = generics[(key[2], key[1])] = dihtype
        generic_type_ids.add(id(dihtype))
    else:
        if id(dihtype) in specific_type_ids:
            continue
        specifics[key] = dihtype
        specific_type_ids.add(id(dihtype))

# Now we have a separation of specifics and generics
print('Found %d generic torsions and %d specific torsions' %
        (len(generic_type_ids), len(specific_type_ids)))

for specific_key, dihtype in specifics.items():
    # Look through all generic terms and see if there are any generics that
    # match. If there is one (there will be ONLY one) see that the specific
    # torsion overrides all periodicities on the generic torsion.
    genkey = (specific_key[1], specific_key[2])
    if genkey not in generics:
        continue
    gdihtype = generics[genkey]
    genper = {int(x.per) for x in gdihtype}
    speper = {int(x.per) for x in dihtype}
    diff = genper - speper
    if len(diff) > 0:
        print('%-2s-%-2s-%-2s-%-2s is missing overriding periodicities %s' %
                (specific_key[0], specific_key[1], specific_key[2],
                    specific_key[3], ', '.join(str(x) for x in sorted(diff)))
        )
```
